# In situ programming of intratumoural stem cell-like memory CD8⁺ T cells enables durable antitumour immunity in immunosuppressive tumours

**DOI:** 10.64898/2026.02.06.700840

**Authors:** Xiaomeng Wang, Ana Vitlic, Erminia Romano, Rachel Scholey, Leo Zeef, Conrado Guerrero Quiles, Antonia Banyard, Richard Walshaw, Tim Illidge, Jamie Honeychurch

## Abstract

Tumours with an immunosuppressive microenvironment often respond poorly to radiotherapy (RT) and immune checkpoint inhibitors, and durable clinical responses remain rare. These failures are not solely explained by insufficient immune activation, but instead reflect an inability of current therapies to establish CD8⁺ T-cell infiltration and states required for long-term tumour control. Although stem cell-like memory CD8⁺ T cells (Tscm) are critical for durable antitumour immunity, they are typically rare within solid tumours, and strategies to selectively augment the expansion of intratumoural Tscm within immunosuppressive tumours are lacking.

Here, we show that intratumoural immune modulation via pattern recognition receptor (PRR) activation, using a double-stranded RNA–based agonist (BO-112), promotes Tscm expansion and, in combination with radiation, enhances antitumour immunity in poorly immunogenic lung and bladder cancer models. PRR activation enabled selective in situ expansion of intratumoural Tscm, rather than global CD8⁺ T-cell activation. Therapeutic efficacy required pre-existing intratumoural CD8⁺ T cells and was associated with qualitative reprogramming of the immune landscape, including depletion of exhausted CD8⁺ T cells and regulatory T cells. Transcriptomic analyses showed that PRR activation preferentially enriched programmes associated with Tscm formation and self-renewal, whereas durable effector differentiation emerged only following combined RT–PRR agonist treatment, consistent with antigen-driven execution of the expanded progenitor pool. A Tscm-associated gene signature derived from these preclinical datasets correlated with improved survival across independent human lung and bladder cancer cohorts. Together, our data identify intratumoural Tscm as a therapeutically targetable immune-control axis and establish in situ programming of Tscm as a route to durable antitumour immunity.

## Introduction

Immune checkpoint inhibitors (ICIs) have transformed cancer treatment, improving clinical outcomes and establishing immunotherapy as a mainstay of cancer management^1^. However, the majority of patients either fail to respond or develop resistance, and long-term tumour control remains uncommon across many solid tumour types^2^. Radiotherapy (RT), which is delivered to over half of all cancer patients^3^, has therefore been widely combined with ICIs to enhance tumour immunogenicity and improve clinical outcomes^3–7^. RT can induce immunogenic tumour cell death, promote tumour antigen release and engage innate sensing pathways that promote dendritic cell (DC) activation and cross-priming of tumour-specific T cells, providing a strong rationale for combination with immunotherapy^5–8^. Preclinical studies have shown that RT can enhance CD8^+^ T-cell-mediated responses and elicit systemic immune effects, including abscopal tumour regression^9–11^. However, translation of RT–ICI combinations has yielded inconsistent and often disappointing results in the clinic, particularly in tumours with an immunosuppressive tumour microenvironment (TME)^8, 12, 13^. These observations highlight a persistent and unresolved barrier to achieving durable antitumour immunity in resistant disease settings.

Recent studies have identified TCF1⁺ stem cell-like memory CD8⁺ T cells (Tscm) as a critical progenitor population capable of sustaining durable antitumour immunity^14–16^. Tscm reside within the memory lineage, possess robust self-renewal capacity, and retain the ability to differentiate into both long-lived memory and effector CD8⁺ T-cell populations required for sustained tumour control^14, 15, 17, 18^. In cancer and chronic infection settings, the presence of Tscm has been associated with durable immune responsiveness and long-term therapeutic benefit^16, 18, 19^. However, Tscm are typically rare within established solid tumours, particularly in immunosuppressive tumour microenvironments^20^. Current immunotherapies, including ICIs and RT-based combinations, drive effector differentiation or expansion of exhausted CD8⁺ T-cell states^21, 22^ and often fail to effectively generate or maintain an intratumoural Tscm pool, potentially explaining the transient nature of many clinical responses. Although adoptive cell therapy (ACT) can generate Tscm ex vivo^14, 15^, this approach is is technically complex, resource-intensive, and not broadly applicable across tumour types^23, 24^, underscoring the need for alternative strategies capable of inducing Tscm in situ.

Pattern recognition receptors (PRRs) detect pathogen– and damage-associated molecular patterns and coordinate innate and adaptive immune responses^25^. Activation of PRRs initiates antiviral immune signalling cascades, promotes antigen presentation, and shapes downstream T-cell differentiation^26,27–30^. These properties suggest that PRR agonism may have the capacity to modulate immune state quality, rather than simply increasing immune activation. Synthetic double-stranded RNA (dsRNA)–based PRR agonists, including those delivered intratumourally, have shown potent immunostimulatory effects in preclinical models and are under clinical investigation^31^. Whether PRR activation can be leveraged to reprogramme intratumoural CD8⁺ T-cell states and access progenitor immunity, however, remains unknown.

Critically, it remains unclear whether intratumoural Tscm can be selectively expanded in situ, particularly within immunosuppressive tumours that respond poorly to RT and ICIs. It is also unknown whether such immune-state modulation can be achieved without systemic immune activation or adoptive cell transfer. Here, we demonstrate that intratumoural PRR activation, using the dsRNA agonist BO-112 in combination with RT, selectively expands Tscm in situ and reprogrammes the intratumoural immune landscape to support durable antitumour immunity in poorly immunogenic tumour models.

## Results

### Intratumoural BO-112 therapy restores sensitivity to RT in radioresistant murine tumour models

To determine whether intratumoural PRR agonism can improve responses in radioresistant tumours, we evaluated the nanoplexed dsRNA agonist BO-112 in combination with hypofractionated RT (3 × 8 Gy) in two murine cancer models: KP475 non–small cell lung cancer (NSCLC; *Kras*^G12D^, *Trp53*^−/−^), and MB49 bladder cancer (Fig. 1A). Consistent with treatment resistance observed in human NSCLC harbouring these oncogenic alterations^32, 33^, KP475 tumours were refractory to RT alone, exhibiting no measurable therapeutic benefit (Fig. 1B). Combination of RT with anti–PD-1 likewise failed to induce tumour control in this model (Supplementary Fig. S1A and B).

**Figure 1.**
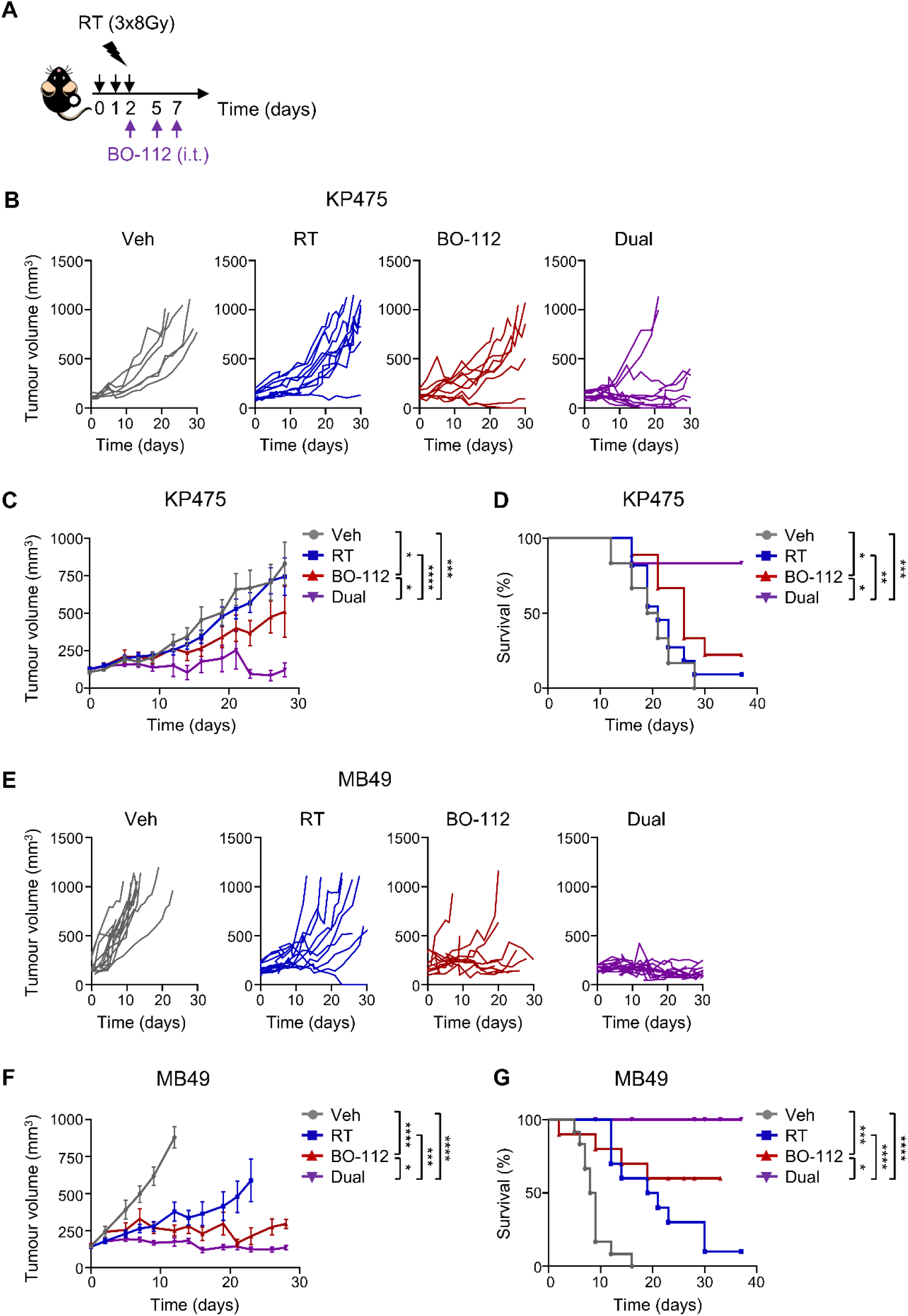
Intratumoural BO-112 therapy restores sensitivity to RT in radioresistant murine tumour models. **A**, Experimental schematic showing that mice were inoculated subcutaneously with KP475 (NSCLC) or MB49 (bladder cancer) cells and treated with RT (3 x 8 Gy), BO-112 (intratumoural injection; i.t.) or their combination (RT + BO-112; Dual). Arrows indicate the treatment start date relative to the first dose of RT. **B,** Individual tumour growth curves, **C,** Tumour volumes (mean ± SEM), and **D,** Kaplan–Meier survival curves for KP475-bearing mice treated with 5% dextrose vehicle (Veh, n = 6), RT + Veh (RT, n = 11), BO-112 (n = 9), or Dual (n = 12). Tumours ranged from 70–200 mm^3^ at treatment start. **E,** Individual tumour growth curves, **F,** Tumour volumes (mean ± SEM), and **G,** Kaplan–Meier survival curves for MB49-bearing mice treated with Veh (n = 12), RT (n = 10), BO-112 (n = 10), or Dual (n = 12). Tumours ranged from 83–200 mm³ at treatment start. All tumour volume and survival data are presented at the indicated time points relative to the first dose of RT. Tumour growth data were analysed using two-way ANOVA (mixed-effects model); survival data were analysed using Log-rank (Mantel-Cox) test. In all cases, * *p* < 0.05, ** *p* < 0.01, *** *p* < 0.001, **** *p* < 0.0001.

While intratumoural administration of BO-112 alone was insufficient to mediate tumour control, its combination with RT resulted in significant tumour growth inhibition and prolonged survival, with sustained tumour control observed in over 80% of treated mice (Fig. 1B–D). Mice exhibiting durable responses rejected subsequent tumour re-challenge, indicating the establishment of tumour-specific immune memory (Supplementary Fig. S1C–E). A similar enhancement of RT efficacy following addition of BO-112 was observed in the MB49 bladder cancer model (Fig. 1E–G). Together, these results show that intratumoural BO-112 therapy restores sensitivity to RT and enables durable tumour control in murine tumours refractory to RT or RT–ICI combination therapy.

### Pre-existing intratumoural CD8^+^ T cells are essential for the therapeutic efficacy of combined RT and BO-112 therapy

We next sought to determine whether the therapeutic efficacy of combined RT and BO-112 was mediated by local or systemic immune responses. To assess the contribution of major T-cell subsets, combination therapy was administered together with antibody-mediated depletion of either CD8^+^ or CD4^+^ T cells (Supplementary Fig. S2A and B). Depletion of CD8^+^ T cell significantly reduced the therapeutic benefit of RT and BO-112 in both the KP475 (Fig. 2A and B; Supplementary Fig. S2C) and MB49 (Fig. 2C and D; Supplementary Fig. S2D) tumour models, confirming that efficacy is CD8⁺ T-cells dependent. In contrast, CD4^+^ T-cell depletion had no significant impact on therapeutic outcome in either model, although a modest reduction in growth control was observed in MB49 tumours (Fig. 2A–D; Supplementary Fig. S2C and D).

**Figure 2.**
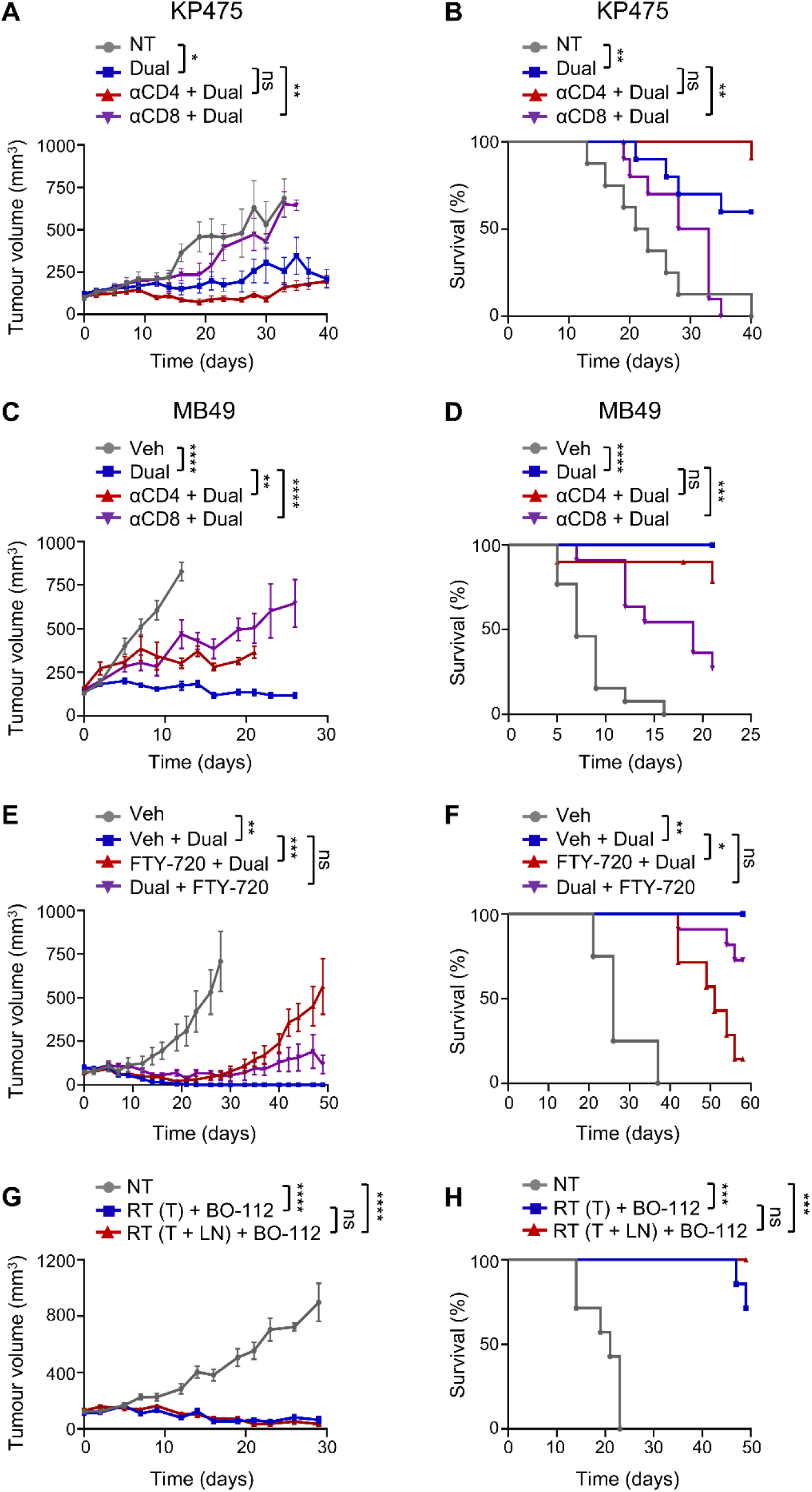
Pre-existing intratumoural CD8^+^ T cells are essential for the therapeutic efficacy of combined RT and BO-112 therapy. **A**, Tumour volumes (mean ± SEM) and **B,** Kaplan–Meier survival curves for mice bearing KP475 tumours treated with no therapy (NT, n = 8), or RT + BO-112 (Dual, n = 10), together with CD4-depleting antibody (αCD4 + Dual, n = 12) or CD8-depleting antibody (αCD8 + Dual, n = 10). Tumour volumes ranged from 50–180 mm^3^ at the initiation of RT. **C,** Tumour volumes (mean ± SEM) and **D,** Kaplan–Meier survival curves for mice bearing MB49 tumours treated with 5% dextrose vehicle (Veh, n = 13), Dual (n = 12), αCD4 + Dual (n = 10) or αCD8 + Dual (n = 11). Tumour volumes ranged from 83–202 mm^3^ at the initiation of RT. **E,** Tumour volumes (mean ± SEM) and **F,** Kaplan–Meier survival curves for mice bearing KP475 tumours treated with HPMC vehicle (Veh, n = 4), Veh + Dual (n = 4), FTY-720 + Dual (FTY-720 starting one day prior to tumour implantation, n = 7) or Dual + FTY-720 (FTY-720 starting one day before the first dose of RT, n = 11). Tumours ranged from 46–173 mm^3^ at the initiation of RT. **G,** Tumour volumes (mean ± SEM) and **H,** Kaplan–Meier survival curves for mice bearing KP475 tumours treated with NT (n = 7), BO-112 in combination with RT delivered to either the tumour alone [RT (T) + BO-112, n = 7] or RT delivered to both the tumour and tumour-draining lymph node [RT (T+LN) + BO-112, n = 6]. Tumour volumes ranged from 60–188 mm^3^ at the initiation of RT. All tumour volume and survival data are presented at the indicated time points following initiation of RT. Tumour growth data were analysed using two-way ANOVA (mixed-effects model) and survival data were analysed using Log-rank (Mantel-Cox) test. In all cases, ns = not significant, * *p* < 0.05, ** *p* < 0.01, *** *p* < 0.001, **** *p* < 0.0001.

To distinguish whether tumour control was mediated by pre-existing intratumoural or newly recruited CD8^+^ T cells, we exploited the requirement for tumour-draining lymph nodes (TDLNs) in priming de novo antitumour T-cell responses^34^. FTY-720, a sphingosine-1-phosphate receptor antagonist that blocks T-cell egress from secondary lymphoid organs^10^, was administered either before tumour implantation (to prevent establishment of intratumoural T cells) or immediately prior to therapy (to preserve pre-existing intratumoural T cells while preventing recruitment of new T cells) (Supplementary Fig. S2E and F). Administration of FTY-720 before tumour implantation significantly compromised therapeutic efficacy, whereas blocking T-cell trafficking immediately prior to treatment did not impair tumour control (Fig. 2E and F; Supplementary Fig. S2G), indicating that pre-existing intratumoural T cells are sufficient to mediate response.

Consistent with this conclusion, analysis using Kikume green:red photoconvertible reporter mice^35^ revealed no significant migration of CD8^+^ T cell from TDLNs to tumours following combination therapy (Supplementary Fig. S3A–C). Furthermore, inclusion of TDLNs within the RT field—resulting in regional lymphocyte depletion^36^ (Supplementary Fig. S3D)—did not alter tumour control or survival compared with tumour-only irradiation (Fig. 2G and H; Supplementary Fig. S3E). Together, these data show that tumour control following RT and BO-112 combination therapy depends on pre-existing intratumoural CD8^+^ T cells.

### Dual therapy alters the intratumoural immune landscape

Given that RT and BO-112 dual therapy depends on intratumoural CD8^+^ T cells, we next analysed immune cell dynamics within the TME of KP475 (NSCLC) tumours treated with RT and BO-112, either alone or in combination. Early immune profiling at day 6 post-treatment revealed a transient reduction in CD8^+^ T cells in tumours receiving dual therapy (Supplementary Fig. S4A; Supplementary Table S1), however, this population subsequently recovered and expanded by day 12 (Fig. 3A and B; Supplementary Table S1). DC populations were reduced on day 6 following BO-112 treatment, either alone or in combination with RT (Supplementary Fig. S4B), and remained reduced at day 12 relative to controls (Fig. 3A; Supplementary Table S1). Notably, despite this overall reduction, the proportion of cDC1s (CD8^+^ DCs) within the remaining DC compartment was significantly increased in tumours treated with the combination therapy (Fig. 3C). The expansion of CD8^+^ T cells at later time points was accompanied by a significant increase in granzyme B^+^ (GZMB^+^) CD8^+^ T cells (Fig. 3D), consistent with enhanced cytotoxic potential. Immunofluorescence imaging corroborated these findings, revealing increased densities of CD8^+^ and GZMB^+^ cells in tumours treated with dual therapy compared with controls (Fig. 3E).

**Figure 3.**
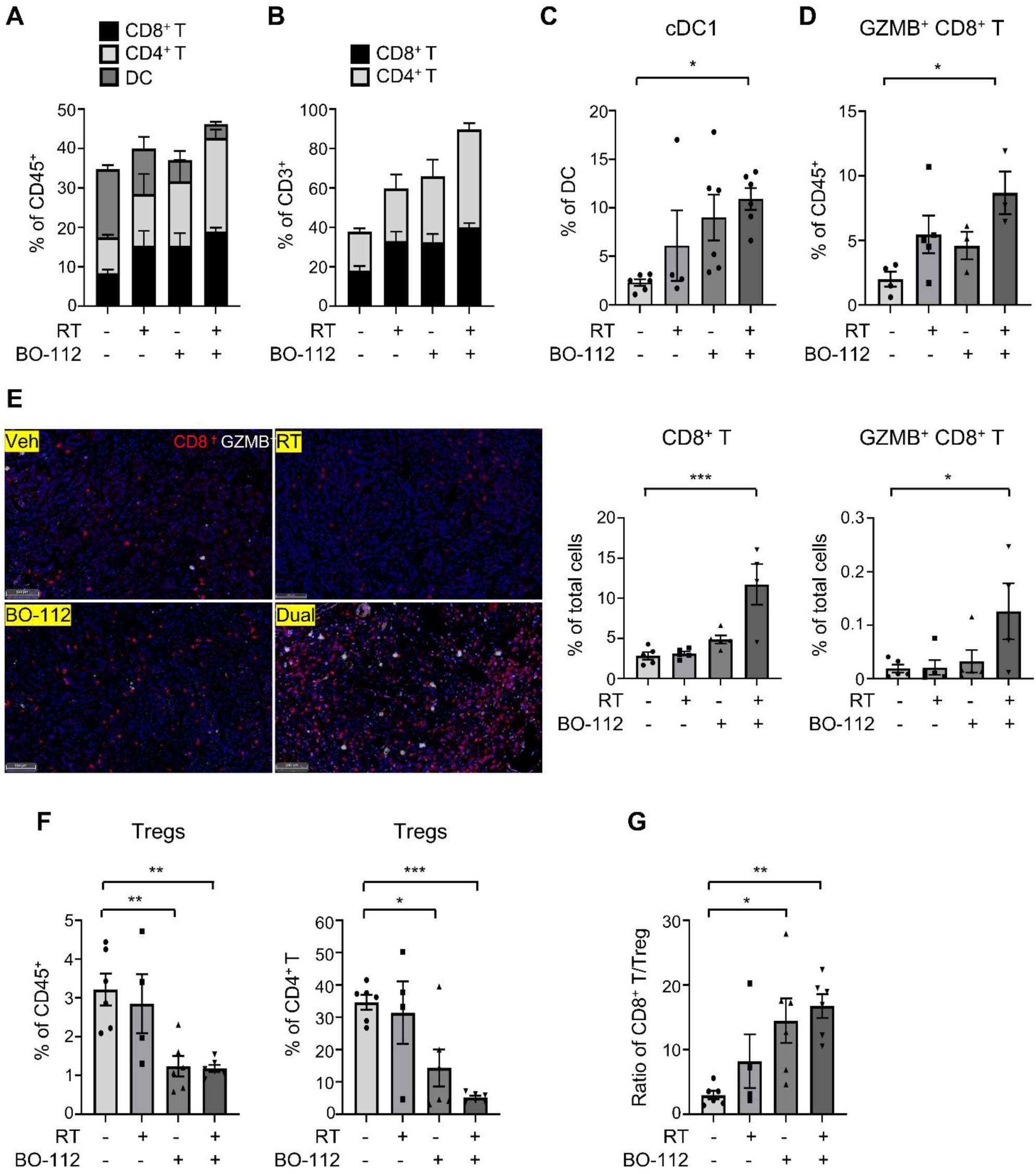
RT and BO-112 dual therapy alters the intratumoural immune landscape. Mice bearing subcutaneous KP475 (NSCLC) tumours were treated with 5% dextrose vehicle (RT–, BO-112–), RT + 5% dextrose (RT+, BO-112–), BO-112 (RT–, BO-112+) or RT + BO-112 dual therapy (RT+, BO-112+). Tumours were surgically harvested on day 12 following RT initiation for downstream analysis. Tumours were either digested into single-cell suspensions for flow cytometry (**A–D**, **F–G**) or fixed in formalin for IHC (**E**). **A,** Frequencies of intratumoural CD8⁺ T cells (CD45⁺ CD3⁺ CD8⁺), CD4⁺ T cells (CD45⁺ CD3⁺ CD4⁺), and DCs (CD45⁺ CD11c⁺ MHC II⁺) within the CD45⁺ immune cell compartment. **B,** Frequencies of CD8⁺ and CD4⁺ T cells within the CD3^+^ lymphocyte population. **C,** Proportion of CD8⁺ DCs (cDC1) among total DCs. **D,** Proportion of granzyme B^+^ (GZMB⁺) CD8⁺ T cells within the CD45⁺ immune compartment. **E,** Representative IHC images and quantification of CD8⁺ T cells (red) and granzyme B (GZMB; white) expression in KP475 tumours across treatment groups. Scale bars, 100 μm. Bar graphs show the proportion of CD8⁺ T cells (left) and GZMB⁺ CD8⁺ T cells (right) among total nucleated cells. **F,** Frequency of Tregs (CD45⁺ CD3⁺ CD4⁺ CD25^+^ Foxp3⁺) within the total CD45⁺ compartment (left) and within the CD4⁺ T cell subset (right). **G,** Ratio of intratumoural CD8⁺ T cells to Tregs. Data are presented as mean ± SEM; each dot represents an individual mouse: n = 3–6 per group as indicated. Statistical analysis was performed using one-way ANOVA with Dunnett’s multiple comparisons test. In all cases, \**p* < 0.05, \*\**p* < 0.01, \*\*\**p* < 0.001, \*\*\*\**p* < 0.0001.

Although total CD4^+^ T-cell populations were increased in tumours receiving combination therapy (Fig. 3A and B; Supplementary Table S1), there was a significant reduction in regulatory T cells (Tregs) within both the CD4^+^ T-cell and total CD45^+^ immune compartments (Fig. 3F). This resulted in a significantly increased CD8^+^ T-cell–to–Treg ratio (Fig. 3G). Selective Treg reduction was also observed following BO-112 monotherapy (Fig. 3F and G) and contrasted with expansion of non-Treg CD4^+^ T cells, indicating selective modulation of CD4⁺ T-cell subsets rather than global depletion. Together, these data show qualitative changes in the intratumoural immune landscape following RT and BO-112 combination therapy, characterised by increased representation of cytotoxic CD8^+^ T-cell subsets and reduced immunosuppressive features.

### Dual therapy enriches stem cell-like memory–associated transcriptomic programmes in intratumoural CD8^+^ T cells

To further elucidate the impact of BO-112 and RT on intratumoural CD8^+^ T cells, we performed RNA sequencing on FACS-isolated CD8^+^ T cells from KP475 tumours treated with vehicle, monotherapies or combination therapy. CD8^+^ T cells were isolated using a defined gating strategy (Supplementary Fig. S5A). Transcriptomic analysis revealed distinct gene expression programmes across treatment groups, with BO-112 emerging as the dominant driver of CD8^+^ T-cells state reprogramming (Fig. 4).

**Figure 4.**
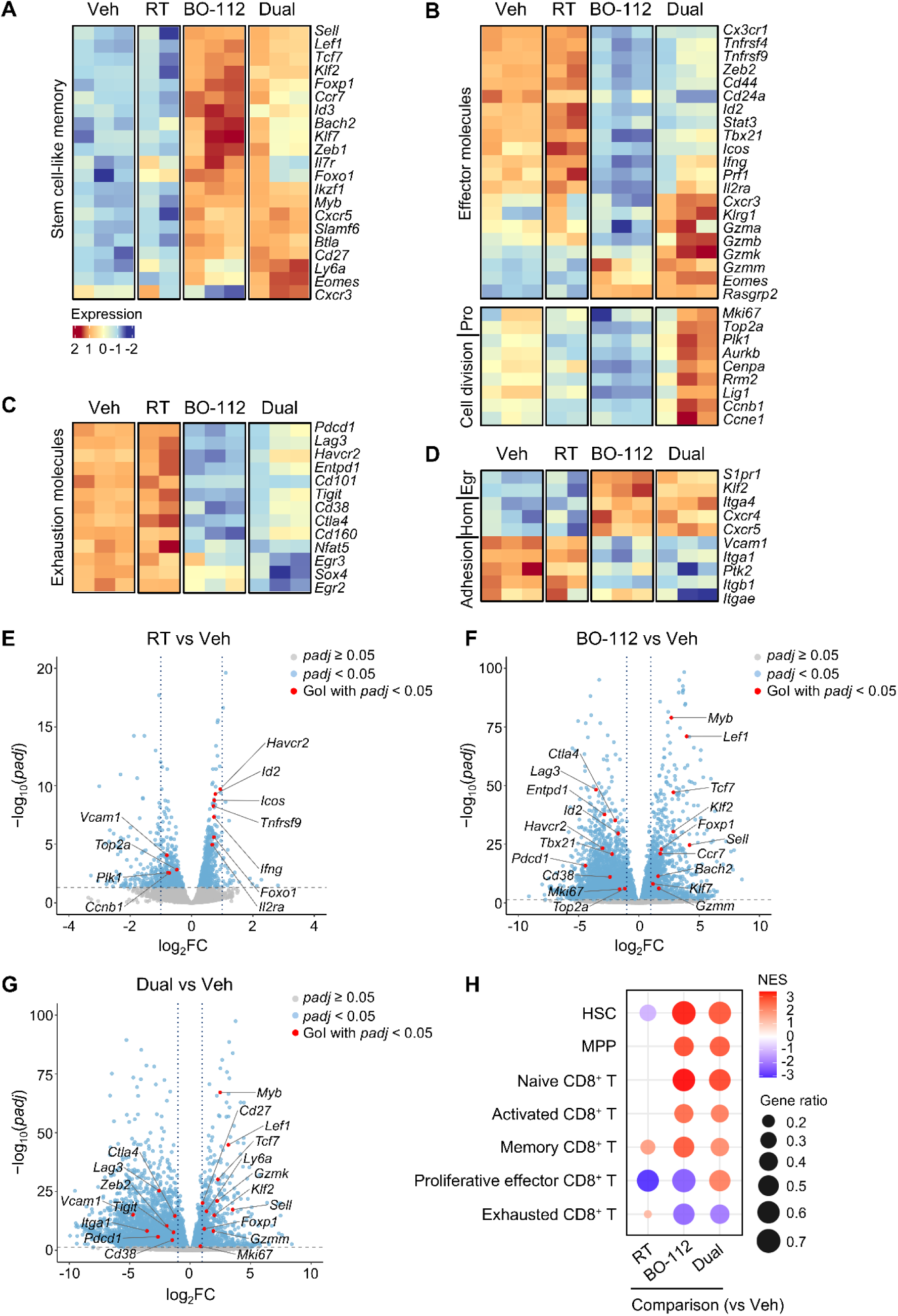
Dual therapy enriches stem cell-like memory–associated transcriptomic programmes in intratumoural CD8^+^ T cells. Mice were implanted with KP475 (NSCLC) tumours and treated with 5% dextrose vehicle (Veh), RT + Veh (RT), BO-112 or RT + BO-112 (Dual). Live CD8^+^ intratumoural lymphocytes were isolated by FACS from tumours on day 12 following the first RT fraction, and bulk RNA was extracted for sequencing. Each sample was pooled from 3–7 tumours. Data shown represent *n* = 2–3 pooled RNA samples per group. **A**–**D,** Heatmaps showing the expression of genes associated with **A,** stem cell-like memory; **B,** effector function, cell division and proliferation (Pro); **C,** exhaustion; and **D,** tissue positioning programmes, including adhesion, niche-homing (Hom), and egress (Egr). Expression values were log_2_ transformed and row-scaled (z-score normalised per gene), ranging from –2 (blue) to 2 (red), indicating relative expression across conditions. **E**–**G,** Volcano plots showing differential gene expression for **E,** RT vs Veh; **F,** BO-112 vs Veh; and **G,** Dual vs Veh comparisons. The x-axis shows log_2_ fold change (log_2_FC), and the y-axis shows –log_10_(*padj*). Grey dots: genes with *padj* ≥ 0.05; blue dots: genes with *padj* < 0.05; red dots: selected genes of interest (GOI) with *padj* < 0.05. **H,** Bubble plot of GSEA using the C7 immunological gene signature collection on CD8^+^ T cell RNA-seq data. Gene sets include haematopoietic stem cells (HSC), multipotent progenitor cells (MPP), naive, activated, memory, proliferative effector, and exhausted CD8^+^ T cells. Comparisons shown: RT vs Veh, BO-112 vs Veh, Dual vs Veh. Normalized enrichment score (NES) ranges from –3 (blue) to 3 (red). Bubble size represents gene ratio (0.2–0.7). Only gene sets with *padj* < 0.25 are shown.

CD8^+^ T cells from BO-112–treated tumours exhibited a transcriptional programme characteristic of stem cell-like memory T cell (Tscm), marked by increased expression of transcription factors associated with memory lineage commitment and self-renewal (*Tcf7*, *Lef1*, *Myb*, *Id3*, *Zeb1*, *Bach2*, *Foxp1*, *Foxo1*, *Klf2* and *Klf7*), surface markers (*Sell*, *Il7r* and *Cd27*), and chemokine receptors (*Ccr7* and *Cxcr5*)^19, 37^ (Fig. 4A and F; Supplementary Fig. S5B). Addition of RT to BO-112 further increased expression of genes associated with activated or transitional Tscm states, including *Ly6a*, *Eomes* and *Cxcr3*^38, 39^, while modestly reducing expression of several early Tscm-associated markers (*Foxp1*, *Ccr7*, *Id3*, *Bach2*, *Il7r* and *Foxo1*) (Fig. 4A and G; Supplementary Fig. S5C). Despite this activation overlay, the overall Tscm-associated transcriptional programme remained dominant following BO-112 + RT treatment (Fig. 4A and G). In contrast, RT alone had minimal impact on Tscm-associated gene expression (Fig. 4A and E), reinforcing BO-112 as the primary driver of the Tscm programme.

Although enrichment of CD8^+^ Tscm states is associated with sustained immune competence^37^, BO-112 monotherapy was insufficient to mediate durable tumour control in vivo. We therefore examined additional transcriptional programmes associated specifically with the dual-therapy context. BO-112 alone suppressed programmes linked to effector differentiation (*Cx3cr1*, *Tnfrsf4*, *Tnfrsf9, Zeb2*, *CD44, Id2, Stat3, Tbx21*, *Icos, Ifng, Prf1* and *Il2ra*)^40, 41^, proliferation (*Mki67* and *Top2a*)^41^ and cell-cycle progression (*Aurkb*, and *Lig1*)^41^ (Fig. 4B and F), consistent with maintenance of a progenitor-enriched state. In contrast, addition of RT induced cytotoxic and proliferative programmes, including effector molecules (*Gzma*, *Gzmb, Gzmk* and *Klrg1*) and cell-division regulators (*Mki67, Top2a*, *Plk1*, *Aurkb*, *Rrm2*, *Ccnb1* and *Ccne1*) (Fig. 4B and G; Supplementary Fig. S5C). These changes are compatible with antigen-driven execution of the expanded progenitor pool and align with the durable tumour control observed following dual therapy, but not BO-112 monotherapy.

Across treatment groups, expression of multiple exhaustion-associated genes, including *Entpd1*, *Cd101*, *Cd38*, *Ctla4*, *Tigit*, *Lag3*, *Pdcd1* and *Havcr2*^40^ was significantly reduced following BO-112 treatment alone or in combination with RT (Fig. 4C, F and G), indicating a shift away from dysfunctional CD8⁺ T-cell states. Gene set enrichment analysis (GSEA) using the MSigDB C7 (Immunologic Signatures) collection confirmed enrichment of Tscm/naive-like CD8^+^ T cell gene sets, as well as haematopoietic stem cell (HSC) and multipotent progenitor (MPP) signatures following BO-112 treatment, which were retained in the dual-therapy setting alongside additional enrichment of activated and proliferative effector CD8^+^ T-cell signatures (Fig. 4H). In contrast, RT alone showed limited enrichment of stem cell-associated programmes. CIBERSORTx deconvolution analysis revealed similar trends, with BO-112 and dual therapy increasing the proportion of naive-like CD8⁺ T cells and reducing exhausted CD8⁺ T cell populations (Supplementary Fig. S5D). GSEA using the MSigDB C2 (Reactome) collection revealed enrichment of T cell receptor (TCR) signalling and cell cycle checkpoint pathways in the dual therapy group (Supplementary Fig. S5E), further supporting a shift toward effector differentiation and proliferative potential.

In addition to state-associated transcriptional programmes, BO-112 and combination therapy altered expression of genes linked to tissue positioning and cellular mobility. This included increased expression of genes associated with Tscm egress (*S1pr1* and *Klf2*)^42, 43^ and niche-directed homing receptors (*Itga4*, *Cxcr4 and Cxcr5*)^37^, alongside reduced expression of adhesion-related genes (*Vcam1*, *Itga1*, *Ptk2*, *Itgb1* and *Itgae*)^44^ (Fig. 4D, F and G). Consistent with these changes, GSEA using the MSigDB C5 (GO:BP) collection revealed downregulation of integrin-mediated adhesion and extracellular matrix interaction pathways (Supplementary Fig. S5F), supporting a shift away from tissue retention towards increased migratory potential.

Collectively, these transcriptomic analyses show that BO-112 treatment establishes a stem cell-like memory–enriched CD8⁺ T-cell state while suppressing exhaustion-associated programmes, and that combination with RT is associated with context-dependent induction of effector and proliferative transcriptional programmes, together reflecting a qualitatively reprogrammed CD8⁺ T-cell landscape.

### Dual therapy activates innate immune programmes and Wnt/β-catenin-associated stemness pathways

As BO-112 is a synthetic dsRNA analogue, we first examined innate immune–associated transcriptional programmes in intratumoural CD8⁺ T cells. BO-112 treatment led to increased expression of genes encoding RIG-I-like receptors (RLRs), including *Dhx58* (LGP2), *Rigi* (RIG-I) and *Ifih1* (MDA5)^45^, together with induction of interferon (IFN)-stimulated genes such as *Irf7*, *Irf9*, *Isg15*, *Ifit1*, *Ifit3*, *Stat1*, *Stat2* and *Cxcl10*^46^ (Fig. 5A). These transcriptional changes are consistent with engagement of innate antiviral sensing pathways by dsRNA analogues. In line with this, GSEA using the MSigDB C5 (GO:BP) collection revealed enrichment of innate immune–associated processes in BO-112-treated CD8⁺ T cells, including cytoplasmic PRR signalling, RIG-I signalling, IFN-related pathways (Fig. 5B).

**Figure 5.**
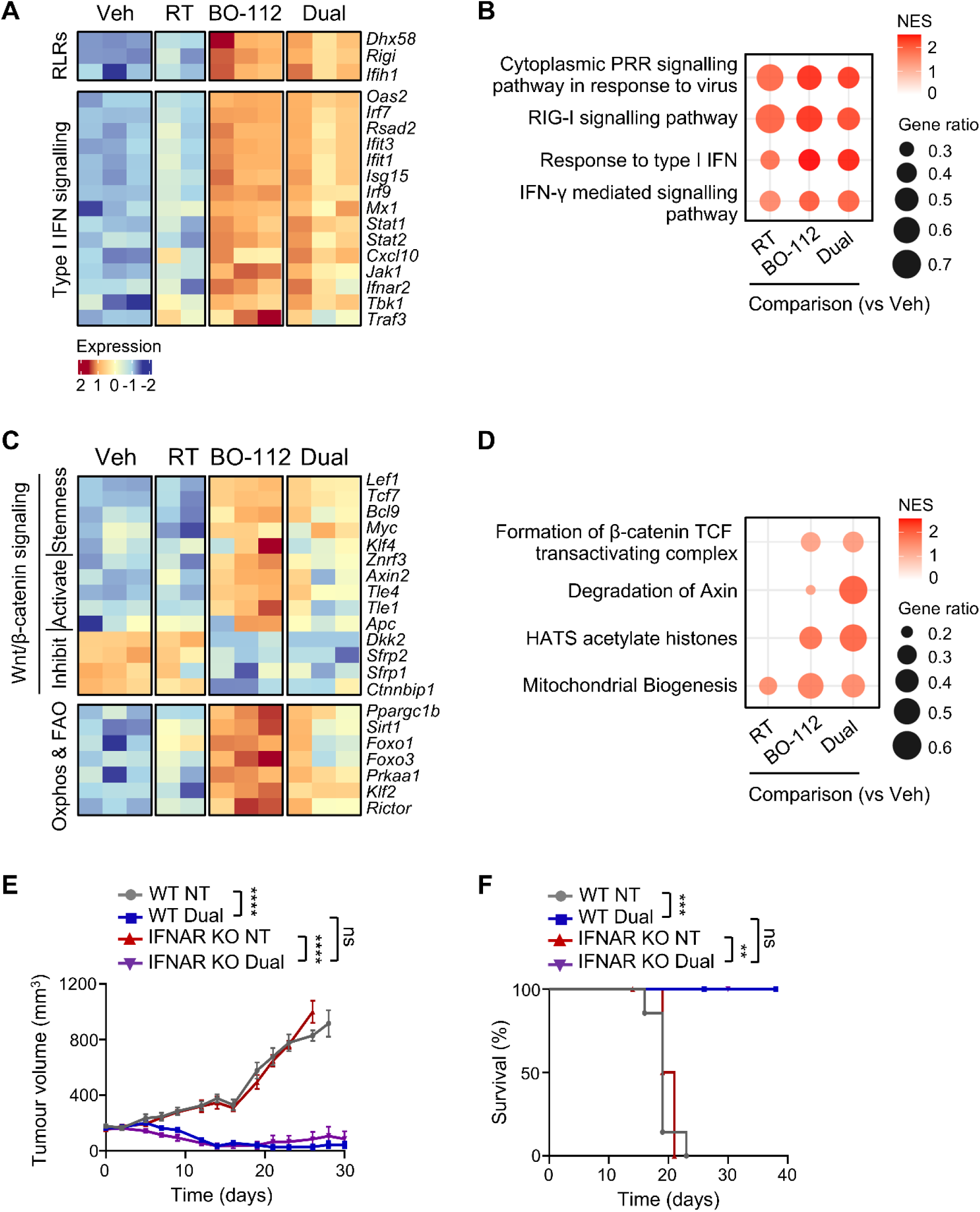
Dual therapy activates innate immune programmes and Wnt/β-catenin-associated stemness pathways. Mice were implanted with KP475 (NSCLC) tumours and treated with 5% dextrose vehicle (Veh), RT + Veh (RT), BO-112 or RT + BO-112 (Dual). Live CD8^+^ T cells were isolated by FACS from tumours on day 12 following the first RT fraction, and bulk RNA was extracted for sequencing. Each sample was pooled from 3–7 tumours. Data represent n = 2–3 pooled RNA samples per group. **A,** Heatmap of genes associated with RIG-I-like receptors (RLRs) and type I IFN signalling, including canonical RLR sensors (e.g., *Dhx58*, *Rigi*) and downstream interferon-stimulated genes (e.g., *Ifit1*, *Isg15*, *Stat1*). **B,** Bubble plot of GSEA using the MSigDB C5 collection (Biological processes), showing enrichment of innate immune pathways, including cytoplasmic PRR signalling and type I IFN response, and IFN-γ signalling. NES ranges from 0 (white) to 2 (red); bubble size represents gene ratio (0.3–0.7). All shown pathways have *padj* < 0.25. **C,** Heatmap of genes associated with WNT/β-catenin signalling and FAO/OXPHOS metabolism. WNT signalling includes stemness-associated transcription factors (e.g., *Tcf7*, *Lef1*, *Myc*) and regulators (*Axin2*, *Dkk2*). Metabolic genes include *Ppargc1b*, *Foxo1*, and *Prkaa1*. **D,** Bubble plot of GSEA using the MSigDB C2 (Reactome pathway) collection, showing enrichment of pathways associated with stemness and epigenetic regulation, including β-catenin TCF complex activity, HATS-mediated histone acetylation, and mitochondrial biogenesis. NES ranges from 0 (white) to 2 (red); bubble size represents gene ratio (0.3–0.7). Only pathways with *padj* < 0.25 are shown. **E,** Tumour volumes (mean ± SEM) and **F,** Kaplan–Meier survival curves for wild-type (WT) or IFNAR knockout (IFNAR KO) C57BL/6 mice bearing KP475 tumours and receiving no treatment (NT; WT n = 7, KO n = 5) or RT + BO-112 dual therapy (Dual; WT n = 5, KO n = 6). Tumour volumes ranged from 137-212 mm^3^ at the initiation of RT. Time on the x-axis indicates days following first fraction of RT. Statistical analysis of tumour growth data was performed using two-way ANOVA (mixed-effects model); survival data was analysed using the Log-rank (Mantel–Cox) test. In both cases, ns = not significant, \*\**p* < 0.01, \*\*\**p* < 0.001, \*\*\*\**p* < 0.0001.

In addition, BO-112 and combination therapy were associated with activation of Wnt/β-catenin signalling, a pathway implicated in maintenance of cellular stemness^17^. This included increased expression of transcription factors (*Lef1*, *Tcf7*, *Bcl9*, *Myc*, and *Klf4*), canonical Wnt target genes (*Znrf3* and *Axin2*), and pathway components (*Tle1*, *Tle4* and *Apc*), together with reduced expression of Wnt signalling inhibitors (*Dkk2*, *Sfrp2*, *Sfrp1*, and *Ctnnbip1*)^47, 48^ (Fig. 5C). Consistent with these gene-level observations, Reactome GSEA showed enrichment of pathways regulating β-catenin–TCF transactivating complex formation and Axin degradation, which stabilizes β-catenin (Fig. 5D), following the dual therapy.

These signalling changes were accompanied by enrichment of metabolic programmes associated with long-term T-cell persistence, including oxidative phosphorylation (OXPHOS) and fatty acid oxidation (FAO). BO-112– and dual-treated CD8⁺ T cells showed increased expression of metabolic regulators (*Ppargc1b*, *Prkaa1*, *Sirt1*, *Foxo1* and *Foxo3*), together with *Klf2* and *Rictor*, which have been implicated in supporting metabolic fitness and durability of Tscm cells^49^ (Fig. 5C). In parallel, Reactome GSEA revealed enrichment of histone acetyltransferase–associated pathways and mitochondrial biogenesis in BO-112– and dual-treated CD8⁺ T cells (Fig. 5D). Together, these data show that BO-112–mediated PRR activation is associated with coordinated engagement of innate immune signalling, stemness-associated transcriptional programmes, metabolic adaptation, and epigenetic regulation in CD8^+^ T cells, consistent with maintenance and expansion of the Tscm phenotype.

Given the enrichment of IFN-related transcriptional programmes following dual therapy, we next assessed whether type I IFN signalling was required for therapeutic efficacy. Wild-type (WT) and IFN-α/β receptor-deficient (IFNAR KO) mice bearing KP475 tumours exhibited comparable tumour control and survival following RT + BO-112 treatment (Fig. 5E and F; Supplementary Fig. S6), showing that host type I IFN signalling is not required for antitumour efficacy of the combination therapy.

### Dual therapy alters the intratumoural T-cell landscape

To validate transcriptomic findings, we performed cytometry by time-of-flight (CyTOF) analysis on intratumoural T cells from KP475 tumours. t-SNE visualization revealed an increased representation of CD8^+^ T-cell populations expressing stemness-associated (TCF-1)^50^, and cytotoxic (GZMB) markers, together with a reduction in populations expressing exhaustion-associated markers such as PD-1, following treatment with BO-112 alone or in combination with RT (Fig. 6A). These changes were concordant with the transcriptional profiles observed by RNA sequencing (Fig. 4). Similar shifts were observed within the CD4^+^ T-cell compartment, including increased TCF-1 expression, reduced exhaustion-associated phenotypes, and a decrease in CD4^+^ FOXP3^+^ regulatory T cells (Tregs) following BO-112-based treatments (Fig. 6A).

**Figure 6.**
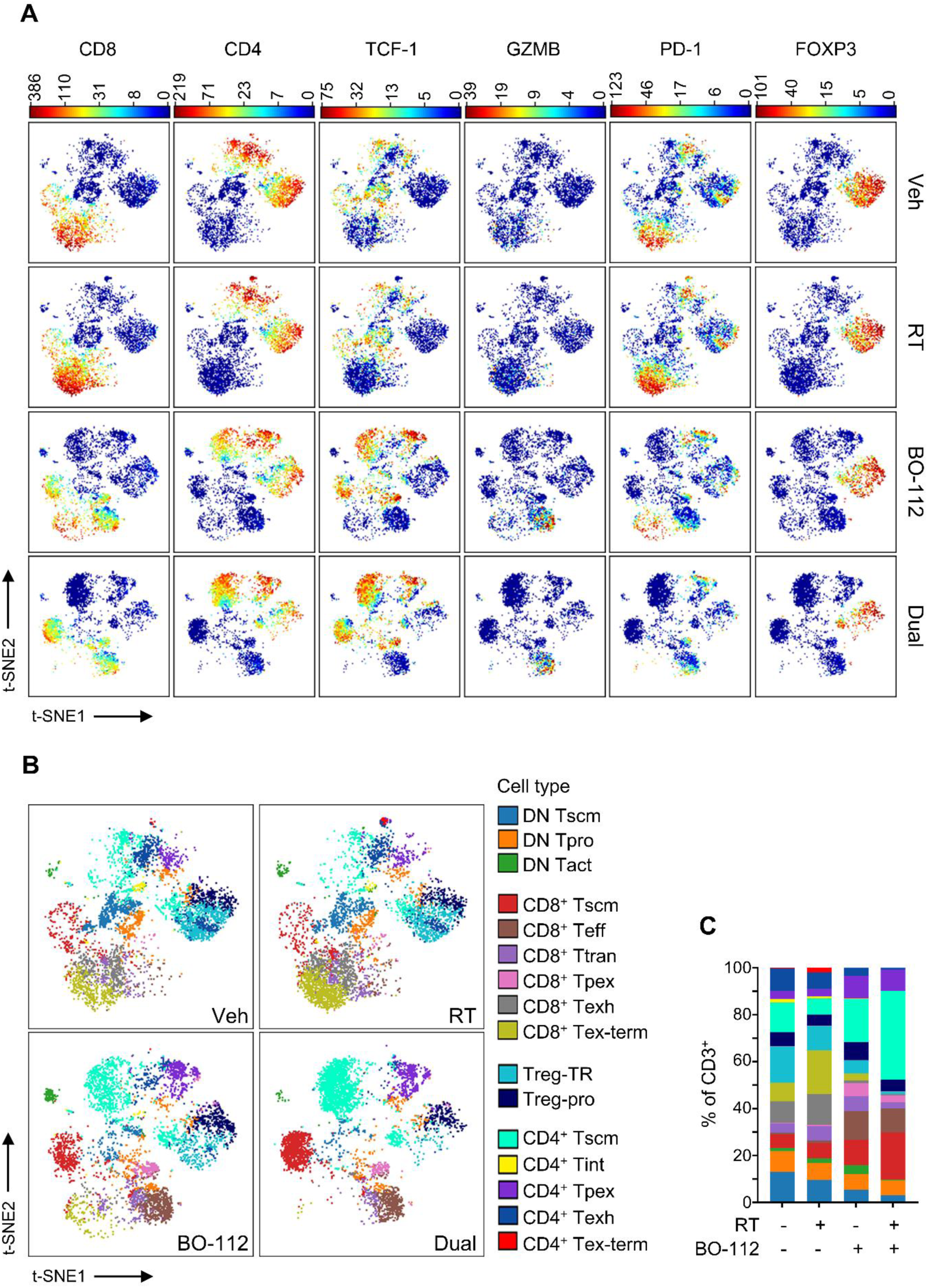
Dual therapy alters the intratumoural T-cell landscape. Mice were implanted with KP475 (NSCLC) tumours and treated with vehicle (Veh), RT + Veh (RT), BO-112 or RT + BO-112 (Dual). Tumours were harvested on day 12 post-first dose of RT, dissociated into single-cell suspensions and pooled per condition (4–5 tumours per group). A total of 3 × 10⁶ cells per group were stained with a panel of metal-conjugated antibodies and analysed by mass cytometry (CyTOF). **A,** Expression of key immune markers (CD8, CD4, TCF-1, GZMB, PD-1, and FOXP3) overlaid on tSNE plots of CD3⁺ T cells isolated from KP475 (NSCLC) tumours under each treatment condition. **B,** FlowSOM clustering and tSNE visualisation of CD3⁺ T cells across treatment conditions. Cell populations were annotated based on marker expression into CD8⁺, CD4⁺, double-negative (DN), and Treg subsets, and further classified into functional states, including stem cell, effector, and exhausted phenotypes. **C,** Relative proportions of each T cell cluster within the total CD3⁺ population per treatment group. Cluster annotations are as follows: Tscm, stem cell-like memory T cells; Tpro, proliferative T cells; Tact, activated T cells; Teff, effector T cells; Ttran, transitory T cells; Tpex, progenitor exhausted T cells; Texh, exhausted T cells; Tex-term, terminally exhausted T cells; Treg-TR, tissue-resident Tregs; Treg-pro, proliferative Tregs; CD4⁺-Tint, intermediate CD4⁺ T cells.

Unsupervised clustering of T cells identified 16 distinct populations encompassing double-negative T cells, CD8^+^ and CD4^+^ stem cell-like memory T cells (Tscm), effector T cells (Teff), precursor exhausted T cells (Tpex), exhausted T cells (Texh), terminally exhausted T cells (Tex-term), and regulatory T-cell subsets (Fig. 6B), defined by marker expression shown in Supplementary Fig. S7. BO-112 treatment resulted in preferential expansion of clusters with phenotypes consistent with stem cell-like memory, characterised by high TCF-1 and CD127 expression with low PD-1 and Ki-67, within both the CD8^+^ and CD4^+^ T-cell compartments. Notably, Tscm constituted a substantially larger fraction of the intratumoural CD8⁺ T-cell compartment than the progenitor-exhausted T cells (Tpex), indicating that Tscm represent the dominant progenitor population following BO-112 treatment. The relative abundance of Tscm was further increased upon addition of RT, such that Tscm represented the dominant subset within both the CD8^+^ and CD4^+^ T-cell pools (Fig. 6C).

In parallel with Tscm enrichment, BO-112 treatment was associated with increased frequency of CD8^+^ T-cell populations exhibiting features of activated effector states, including elevated Ki-67, EOMES, GZMB, PD-1, CD24 and CXCR3, together with intermediate TOX expression, consistent with cytotoxic effector phenotypes (Teff). BO-112 also increased a CD8^+^ T-cell population defined by high Ki-67, TCF-1, and PD-1 expression, consistent with a proliferative, progenitor-like exhaustion-associated state (Tpex) (Fig. 6B and C). However, Tpex consistently represented a markedly smaller population than Tscm across BO-112–based treatment conditions. Although the relative proportions of Teff and Tpex populations were modestly reduced following combination therapy compared with BO-112 alone, these subsets remained more abundant than in vehicle– or RT-treated tumours, while Tscm increased disproportionately, consistent with a rebalancing of the CD8⁺ T-cell compartment towards a stem-like memory–enriched state under dual therapy.

Consistent with the transcriptomic analyses, exhausted CD8^+^ T-cell populations—including Texh (intermediate Ki-67, TCF-1 and TOX with high PD-1) and terminally exhausted Tex-term (loss of TCF-1 with high TOX, PD-1, LAG3 and CD38)—were predominant in vehicle– and RT-treated tumours but were markedly reduced following BO-112 treatment, with a further decrease observed upon addition of RT (Fig. 6B and C). Similarly, CD4^+^ tissue-resident Treg (Treg-TR) were reduced following BO-112 treatment, with the greatest reduction observed in the dual-therapy group. Together, these results show that BO-112, particularly in combination with RT, alters the intratumoural T-cell landscape, with preferential enrichment of CD8⁺ and CD4⁺ Tscm and reduction of exhausted and immunosuppressive T-cell subsets.

### Tscm gene signature correlates with improved survival in clinical cancer cohorts

To assess the clinical relevance of our findings, we applied a Tscm gene signature derived from preclinical RNA sequencing of intratumoural CD8⁺ T cells (Fig. 4A; Supplementary Table S2) to publicly available datasets from patients with NSCLC and advanced bladder cancer. Patients were stratified into high and low Tscm signature groups based on the median signature score. In the TCGA-LUAD cohort, Kaplan–Meier survival analysis showed significantly improved overall survival in patients with high Tscm scores (*p* = 0.0017; Fig. 7A). A similar association was observed in the IMvigor210 cohort of advanced urothelial cancer patients treated with anti–PD-L1 therapy (*p* = 0.021), as well as in the GSE72094 LUAD cohort (*p* = 0.023) (Fig. 7A).

**Figure 7.**
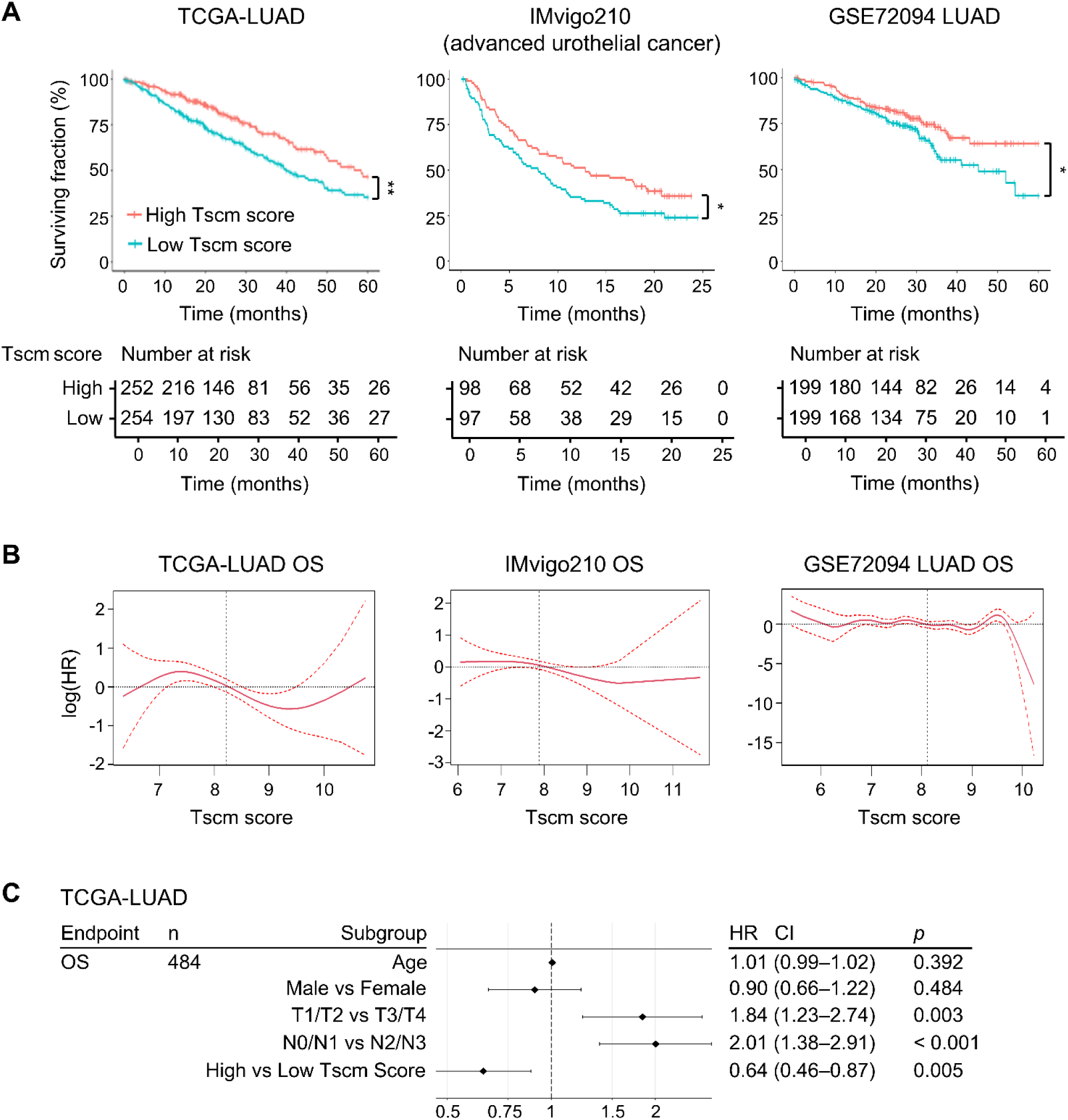
Tscm gene signature correlates with improved survival in clinical cancer cohorts. **A**, Kaplan–Meier survival curves comparing overall survival between patients with high versus low Tscm gene signature scores in the TCGA-LUAD (lung adenocarcinoma; n = 506), IMvigor210 (urothelial carcinoma; n = 195), and GSE72094 (lung adenocarcinoma; n = 398) cohorts. Patients were stratified by median dichotomisation of the Tscm gene signature score, calculated as the median expression *SELL, LEF1, TCF7, KLF2, CCR7, ID3, BACH2, IL7R, FOXO1, IKZF1, CXCR5, SLAMF6, BTLA,* and *CD27*. *P* values were calculated using the log-rank (Mantel–Cox) test: \*\**p* = 0.0017 (TCGA-LUAD), \**p* = 0.021 (IMvigor210), and \**p* = 0.023 (GSE72094). **B,** Cox proportional hazards spline models illustrating the association between the continuous Tscm signature score and overall survival (OS) in the TCGA-LUAD, IMvigor210, and GSE72094 cohorts. Spline fits were generated using penalised smoothing splines within Cox regression models to visualise non-linear relationships between Tscm score and log hazard ratio (log[HR]). The solid grey line represents the estimated log[HR], and the red shaded region denotes the 95% confidence interval. A log[HR] below zero indicates a survival benefit associated with higher Tscm scores. In all cohorts, the confidence intervals widened at the extremes of the Tscm score range due to fewer patients in those ranges. **C,** Forest plot showing the results of multivariable Cox proportional hazards regression analysis in the TCGA-LUAD cohort (n = 484). The model includes clinical covariates (age, sex, tumour stage, and lymph node status) alongside the Tscm signature score (high vs low, median dichotomised). Hazard ratios (HR), 95% confidence intervals (CI), and *p* values are shown for each variable. High Tscm score was independently associated with improved overall survival (HR = 0.64, 95% CI = 0.46–0.87, *p* = 0.005), after adjusting for clinical factors.

To further evaluate the relationship between the Tscm signature and survival, we performed Cox proportional hazards modelling. Treating the Tscm signature score as a continuous variable revealed a graded association with survival. In TCGA-LUAD, increasing Tscm scores were associated with a progressively reduced hazard of death (*p* = 0.0006; Fig. 7B). A similar, though weaker, trend was observed in the IMvigor210 cohort (*p* = 0.06), whereas no consistent association was detected in the GSE72094 LUAD cohort (Fig. 7B). Importantly, multivariate Cox regression analysis in TCGA-LUAD showed that a high Tscm signature remained an independent prognostic factor (HR = 0.64, 95% CI = 0.46–0.87, *p* = 0.005), after adjustment for age, sex, and tumour stage (T and N) (Fig. 7C). Collectively, these data show that enrichment of Tscm-associated transcriptional programmes is associated with improved survival across independent NSCLC and bladder cancer cohorts, consistent with the clinical relevance of progenitor CD8⁺ T-cell states in human cancer.

## Discussion

In this study, we identify intratumoural stem cell-like memory CD8⁺ T cells (Tscm) as a therapeutically targetable immune-control axis within the TME underlying durable antitumour immunity in immunosuppressive tumours. Through intratumoural activation of PRRs in combination with RT, we demonstrate that selective expansion of tumour in situ Tscm is mechanistically linked to therapeutic efficacy in tumour models refractory to RT and ICI. Across transcriptomic, metabolic and cytometric analyses, PRR agonism via BO-112 consistently promoted a CD8^+^ T-cell state characterised by TCF1-associated stemness programmes, self-renewal capacity, and reduced exhaustion, while concurrently depleting Tregs. These changes were not observed with RT alone, indicating that PRR agonism qualitatively reprogrammes intratumoural CD8^+^ T-cell fate rather than amplifying immune magnitude. When combined with RT, this reprogramming supports sustained CD8⁺ T-cell states capable of long-term immune control^22^.

Strategies to expand Tscm within solid tumours are highly desirable but currently lacking. Tscm are defined by a quiescent, multipotent phenotype and the capacity to sustain antitumour immunity through long-term persistence and effector generation^15^. Using CyTOF, we identified two distinct TCF1^+^ CD8⁺ T-cell subsets following BO-112 treatment: a smaller PD-1^high^ Ki67^high^ progenitor-exhausted (Tpex) subset and a dominant PD-1^low^ Ki67^low^ subset consistent with Tscm, which was further expanded by the addition of RT. This phenotypic shift was accompanied by enrichment of Tscm-associated gene signatures and metabolic programmes linked to memory formation and quiescence, including oxidative phosphorylation and fatty acid oxidation^51^. Together, these data support a model in which BO-112 selectively expands a less exhausted, stem-like memory CD8⁺ T-cell compartment in situ. Notably, the abundance of this Tscm population correlated with tumour control in models refractory to RT and ICI, suggesting that intratumoural expansion of Tscm may represent a more effective strategy than attempting to reinvigorate exhausted or numerically limited Tpex populations.

RT is widely used in combination with immunotherapies, yet the precise mechanisms by which it enhances immune efficacy—particularly in synergy with agents like ICIs—have not been fully elucidated^4^. RT is known to promote antigen release^6^ and initiate pro-inflammatory signalling^13^, but whether it enhances antitumour immunity by amplifying immune magnitude, altering T-cell differentiation, or licensing effector function remains an open question^4^. In our study, BO-112 appeared to be the primary driver of immune-state reprogramming, expanding a stem-like TCF1⁺ PD-1^low^ Tscm population. However, this alone was insufficient to mediate durable tumour control. Transcriptomic analysis revealed that BO-112-expanded Tscm did not intrinsically differentiate into cytotoxic effectors. In contrast, the addition of RT led to the emergence of CD8⁺ T cells expressing high levels of granzymes, indicative of cytolytic function. These data further refine our model by revealing a division of labour in which BO-112 establishes a self-renewing, exhaustion-resistant progenitor reservoir, while RT provides the complementary, context-specific signals required to license effector differentiation and drive tumour clearance^6, 13^.

Although PRR agonists have been primarily studied for their ability to stimulate innate immunity and enhance antigen-presenting cell activation, their capacity to programme distinct CD8^+^ T-cell fate within the TME requires further characterisation^25, 26^. In our study, BO-112—a synthetic dsRNA analogue—induced transcriptional programmes associated with T-cell stemness, including Wnt/β-catenin signalling, oxidative phosphorylation, fatty acid oxidation, and epigenetic pathways previously implicated in the persistence of self-renewing memory T cells^52^. These changes coincided with selective expansion of intratumoural TCF1^+^ PD-1^low^ Tscm, and therapeutic efficacy was preserved in IFNAR-deficient hosts, indicating that type I IFN responsiveness in immune cells is dispensable for tumour control. Instead, our data suggest a role for non-canonical outputs downstream of RIG-I and MDA5 in shaping CD8^+^ T-cell fate. Notably, both sensors have been linked to inflammation-induced haematopoietic progenitor generation^53^, pointing to a broader connection between PRR signalling and stemness induction across immune lineages. Whether Tscm expansion following BO-112 treatment is mediated via Wnt/β-catenin remains to be determined. Interestingly, this pathway may also destabilise Treg by suppressing Foxp3 transcription^54^, potentially contributing to the concurrent reduction in Tregs observed with BO-112 treatment. Collectively, our data broaden the functional scope of PRR agonists beyond innate activation, revealing their capacity to reprogramme intratumoural CD8^+^ T-cell states toward stem-like fates.

Anatomical compartmentalisation of immune activation has important implications for immunotherapy efficacy, yet the spatial requirements for effective CD8^+^ T-cell programming remain incompletely understood^55^. Stem-like CD8^+^ T cells—including Tscm and Tpex—can circulate between tumours and TDLNs^56^, and TDLNs have been shown to serve as reservoirs for stem-like progenitors capable of sustaining responses to therapies such as PD-1 blockade^57, 58^. However, the extent to which these peripheral reservoirs are required across distinct treatment modalities remains unclear. In our study, therapeutic efficacy of BO-112 plus RT was preserved even when T-cell trafficking into tumours was blocked prior to treatment, indicating that the response derives predominantly from cells resident in the tumour at the time of therapy. This suggests that in situ expansion and differentiation of intratumoural Tscm is sufficient to mediate tumour control in this setting. Although a contributory role for TDLNs in maintaining Tscm pools during tumour evolution cannot be excluded, these findings contrast with immunotherapies that rely on systemic priming or peripheral reservoirs—such as PD-1/PD-L1 blockade or intratumoural CD40 agonists^59^. Instead, the mechanism of BO-112 plus RT may more closely resemble that of CTLA-4 blockade, where modulation of intratumoural immune populations is central to therapeutic efficacy^60^. Notably, this localised mode of action may offer clinical advantages in contexts where lymph nodal priming is impaired, such as during elective nodal irradiation^61^, and supports the broader potential of in situ immune programming strategies in tumours resistant to systemic ICI therapies.

Despite increasing recognition of CD8^+^ T cell heterogeneity, the clinical relevance of specific transcriptional states—particularly progenitor-like populations such as Tscm—has not been clearly established^62^. To address this, we analysed human NSCLC and bladder cancer datasets for the presence and prognostic significance of a Tscm-associated gene signature derived from our murine RNA-seq data. Across three independent cohorts (TCGA-LUAD, GSE72094, and IMvigor210), high Tscm scores were consistently associated with improved overall survival, supporting the clinical relevance of this immune-state framework. In TCGA-LUAD, the Tscm signature remained an independent prognostic factor after adjusting for tumour stage, indicating that it captures information beyond standard clinical variables. While the IMvigor210 cohort involved anti–PD-L1 treatment, the association with outcome was interpreted as evidence of general immune competence rather than predictive response. Our data suggest that the Tscm programme reflects a favourable intratumoural immune landscape and support the potential of therapeutic strategies—such as BO-112 plus RT—that expand this population in situ.

In conclusion, our study identifies intratumoural Tscm as a therapeutically targetable immune-control axis capable of sustaining durable antitumour immunity. By demonstrating that local PRR agonism selectively expands Tscm and that RT converts this progenitor pool into functional effectors, we establish a mechanistic framework that explains the limitations of effector-centric immunotherapies in immunosuppressive tumours. These findings position immune-state programming—rather than immune activation alone—as a critical determinant of therapeutic success, offering a new design principle for next-generation immunotherapies. Importantly, the clinical relevance of this axis is supported by consistent association between the Tscm gene signature and improved survival across independent human cancer cohorts. Together, these insights argue for a paradigm shift: from amplifying immune magnitude to reprogramming immune cell fate in situ as a strategy to overcome resistance and achieve durable tumour control.

## Materials and Methods

### Cell lines

KP475 murine NSCLC and MB49 murine bladder cancer cells were cultured in F-12/DMEM or DMEM, respectively, supplemented with 10% foetal bovine serum. Cells were maintained at 37 °C with 5% CO₂ and routinely tested for mycoplasma contamination.

### Mouse models

C57BL/6 wild-type mice, Kikume Green:Red photoconvertible mice, and IFNAR knockout mice were used as indicated. Mice were housed under standard specific-pathogen-free conditions and used at 7–12 weeks of age. All animal studies were performed under a United Kingdom Home Office Project license (PCC943F76 and PP1231845) and approved by the CRUK Manchester Institute Animal Welfare and Ethical Review Board (AWERB).

### In vivo treatment

Subcutaneous tumours were established by injecting 1 × 10⁶ KP475 or MB49 cells into the flank of mice. Treatments were initiated when tumours reached ∼75–200 mm^3^. RT was delivered as three daily fractions of 8 Gy (3 × 8Gy) using a cabinet X-ray irradiator with tumour shielding. BO-112 (Highlight Therapeutics) or vehicle control (5% dextrose) was administered intratumourally. Anti-CD4, anti-CD8 and anti-PD-1 antibodies (Bio X Cell) were administered intraperitoneally. FTY-720 (Fingolimod; Enzo Life Sciences) was administered by oral gavage.

### Flow cytometry and immunohistochemistry (IHC)

Tumours were enzymatically dissociated to generate single-cell suspensions for flow cytometric analysis. Cells were stained with viability dyes and fluorochrome-conjugated antibodies (Supplementary Table S3), acquired on a flow cytometer, and analysed using FlowJo software. Formalin-fixed paraffin-embedded tumour sections were stained for CD8 (eBioscience), Granzyme B (Abcam), and γH2AX (CST) using chromogenic or multiplex immunofluorescence approaches and quantified using HALO software.

### CD8⁺ T cell isolation and RNA sequencing

Intratumoural CD8⁺ T cells were isolated by fluorescence-activated cell sorting (FACS) from pooled tumours. Bulk RNA was extracted immediately following sorting, and strand-specific poly(A)-enriched libraries were prepared prior to paired-end sequencing on an Illumina platform. Differential expression analysis was performed using DESeq2, and GSEA was conducted using established MSigDB collections. Differential expression data of genes of interest used for heatmap and volcano plots are provided in Supplementary Table S4.

### Mass cytometry (CyTOF)

Tumour-derived single-cell suspensions were stained with metal-conjugated antibodies (Supplementary Table S5), barcoded, and acquired on a CyTOF XT instrument. Data were normalised, debarcoded, and analysed using tSNE for dimensionality reduction and FlowSOM for clustering.

### Analysis of clinical data

Publicly available transcriptomic and clinical datasets were analysed to assess the clinical relevance of BO-112-associated CD8⁺ Tscm signature. Gene expression scores were calculated using curated Tscm-associated genes (Supplementary Table S2), and associations with survival outcomes were assessed using Kaplan–Meier and Cox regression analyses.

### Statistical Analysis

Statistical analyses were performed using GraphPad Prism 10 software. Tumour growth data were analysed using a mixed-effects model with time and treatment as fixed effects, including a time × treatment interaction. Survival data were analysed using Kaplan–Meier estimates and compared using log-rank (Mantel–Cox) tests. Flow cytometry and IHC data were analysed using one-way ANOVA with Dunnett’s or Tukey’s multiple comparisons tests where appropriate. A *p* < 0.05 was considered statistically significant.

## Data availability

The RNA sequencing data generated in this study are available in the ArrayExpress database under accession number E-MTAB-15281. Transcriptomic and clinical data for the TCGA-LUAD cohort are available from the [Genomic Data Commons (GDC) Data Portal](https://portal.gdc.cancer.gov/projects/TCGA-LUAD) and [cBioPortal](https://www.cbioportal.org/study/summary?id=luad_tcga_pan_can_atlas_2018). Data for the GSE72094 cohort are available in the [Gene Expression Omnibus (GEO)](https://www.ncbi.nlm.nih.gov/geo/query/acc.cgi?acc=GSE72094). The IMVigor210 dataset is accessible via IMvigor210CoreBiologies R package. All remaining data are available within the article, the online supplementary information, or the source data files. Source data are available upon reasonable request.

## Acknowledgements

We thank members of the Cancer Research UK Manchester Institute Core Facilities including the Biological Resources Unit (BRU), led by Lisa Door and her team for their support with in vivo studies; Flow Cytometry (Emily Scanlon and Adam Milner, led by Antonia Banyard); Histology (Caron Behan, led by Garry Ashton); and Molecular Biology (John Weightman and Rachel Horner, led by Wolfgang Breitwieser). We also thank the Christie Hospital Medical Physics for dosimetry and technical support with the radiotherapy platforms. We are grateful to all members of the Targeted Therapy group for helpful discussions. This work was funded by Cancer Research UK(C18915/A29362), FCAECC and AIRC under the Accelerator Award Programme and Cancer Research UK Programme Grant (C431/A28280).

## Author Contributions

X.W. designed the study, performed experiments, analysed data, and wrote the manuscript. A.V. and E.R. designed and performed experiments and analysed data. R.S. and L.Z. developed methodology and analysed data. C.G.Q. analysed data. A.B. performed experiments and data processing. R.W. developed methodology. T.I. acquired funding and supervised the study. J.H. designed and supervised the study and wrote the manuscript. All authors reviewed and edited the manuscript. A.V. and E.R. contributed equally. T.I. and J.H. are joint senior authors.

## Ethics declaration

All animal procedures were performed under a United Kingdom Home Office License (PCC943F76 and PP1231845) approved by the local Animal Welfare and Ethical Review Board (AWERB) at the CRUK-Manchester Institute. All patient transcriptomic and clinical data used in this study were obtained from publicly available repositories (TCGA, GEO, and IMvigor210), for which ethical approval and informed consent were obtained by the original investigators.

## Materials and Correspondence

Corresponding authors: Dr Jamie Honeychurch, and Professor Tim Illidge

