## Supplementary Information for "In situ programming of intratumoural stem cell-like memory CD8⁺ T cells enables durable antitumour immunity in immunosuppressive tumours"

Supplementary Figures

### Supplementary Figure S1

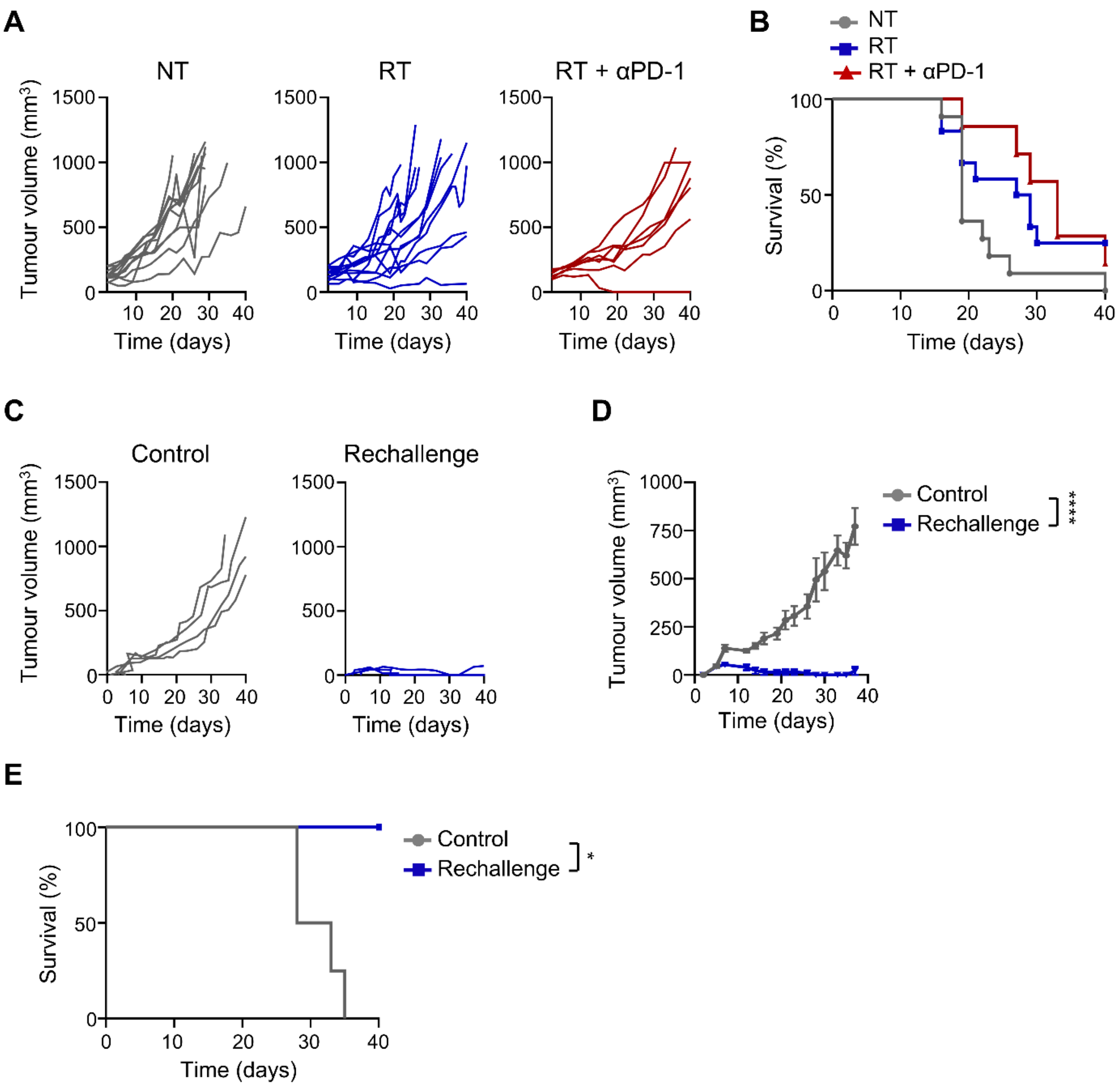

**Supplementary Figure S1.** Synergistic effects of intratumoural BO-112 and RT in KP475 and MB49 tumour models. Related to Figure 1.

**A,** Individual tumour growth curves and **B,** Kaplan–Meier survival curves for mice bearing subcutaneous KP475 NSCLC tumours treated with no therapy (NT, n = 11), RT (n = 12), or RT + αPD-1 (n = 7). Tumours ranged from 97–200 $mm^3$ at treatment initiation. Time indicates days relative to the first RT fraction. **C–E**, Tumour rechallenge in mice that previously rejected KP475 tumours following RT–BO-112 dual therapy. Mice that remained tumour-free for at least 60 days were rechallenged with KP475 cells on the contralateral (left) flank (Rechallenge, n = 3). Control mice (n = 4) were tumour-naive and implanted with KP475 cells for the first time. Time refers to days following rechallenge. **C,** Individual tumour growth curves. **D,** Tumour volumes (mean ± SEM). **E,** Kaplan–Meier survival curves. Tumour growth data were analysed using two-way ANOVA (mixed-effects model) and survival data using Log-rank (Mantel-Cox) test. In all cases, $*p < 0.05$, $****p < 0.0001$.

### Supplementary Figure S2

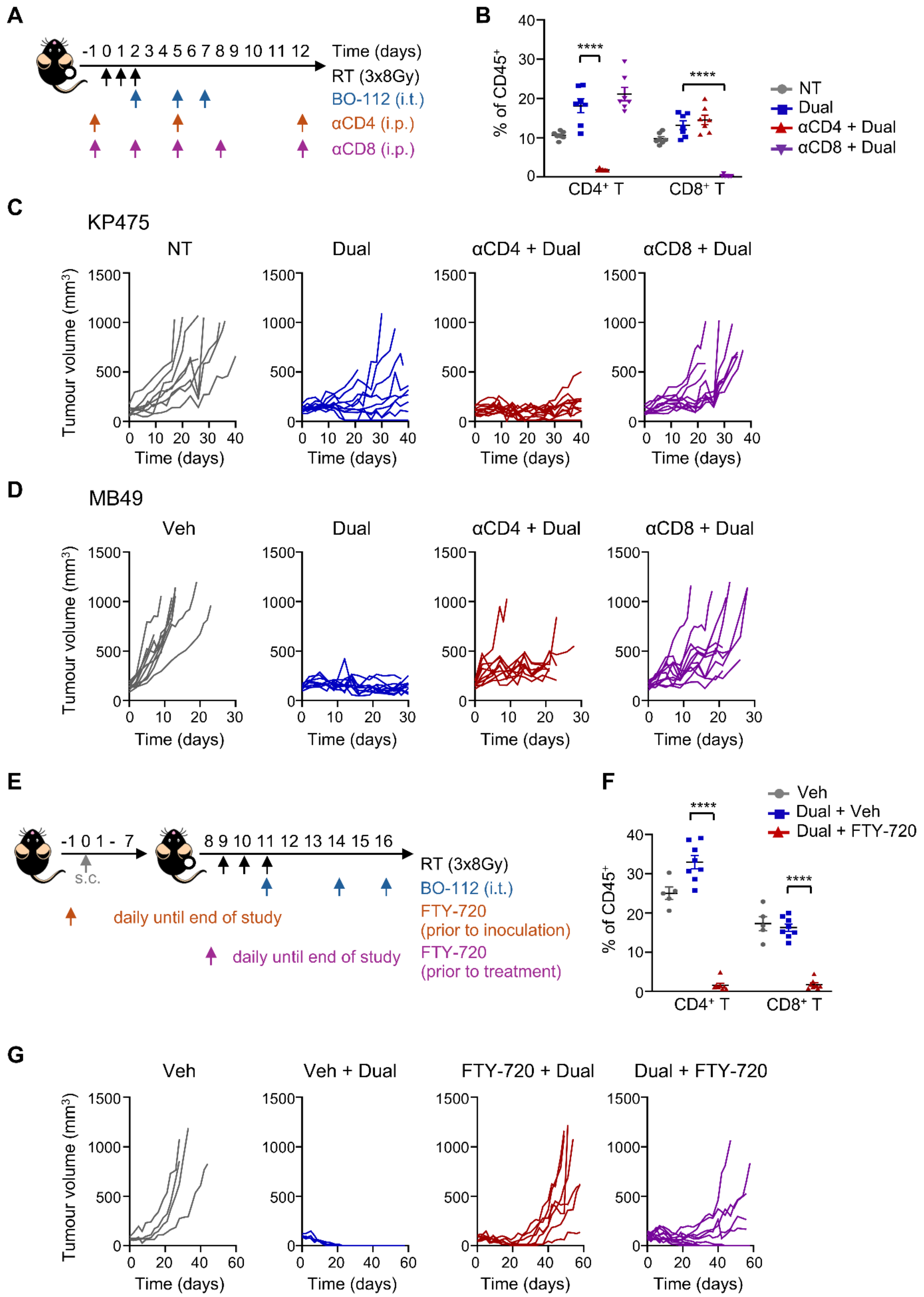

**Supplementary Figure S2.** Tumour in situ $CD8^+$ T cells predominantly mediate the therapeutic efficacy of RT–BO-112 dual therapy. Related to Figure 2.
**A,** Experimental schematic showing that mice were inoculated subcutaneously with KP475 NSCLC or MB49 bladder cancer cells and treated with RT–BO-112 (Dual) in the presence or absence of anti-CD4 or anti-CD8 depleting antibodies. Arrows indicate time relative to the first RT fraction. **B,** Flow cytometry analysis of $CD4^+$ and $CD8^+$ T cell frequencies (mean ± SEM; n = 7 per group) in peripheral blood (tail vein) following antibody-mediated depletion. Each dot represents an individual mouse. **C,** Individual tumour growth curves for KP475 tumour-bearing mice receiving no therapy (NT, n = 8), RT + BO-112 (Dual, n = 10), αCD4 + Dual (n = 12) or αCD8 + Dual (n = 10). Tumour volumes ranged from 50–180 $mm^3$ at RT initiation. **D,** Individual tumour growth curves for MB49 tumour-bearing mice treated with 5% dextrose (Veh, n = 13), Dual (n = 12), αCD4 + Dual (n = 10) or αCD8 + Dual (n = 11). Tumour volumes ranged from 83–202 $mm^3$ at RT initiation. **E,** Mice were inoculated with KP475 cells and treated with Dual therapy (RT + BO-112) in the presence or absence of FTY-720 given either prior to tumour inoculation or prior to treatment. Arrows indicate time relative to tumour implantation. **F,** Flow cytometry analysis of $CD4^+$ and $CD8^+$ T cell frequencies (mean ± SEM) in peripheral blood following FTY-720 administration. Each dot represents an individual mouse: HPMC vehicle (Veh, n = 5), Dual + Veh (n = 8), Dual + FTY-720 (n = 8). **G,** Individual tumour growth curves for KP475 tumour-bearing mice treated with HPMC (Veh, n = 4), Veh + Dual (n = 4), FTY-720 + Dual (FTY-720 initiated one day prior to tumour implantation, n = 7) or Dual + FTY-720 (FTY-720 initiated one day before the first RT fraction, n = 11). Tumour volumes ranged from 46–173 $mm^3$ at RT initiation. All tumour volume measurements are shown at the indicated time points relative to the first RT fraction. One-way ANOVA with Tukey's multiple comparisons test was used for statistic comparison. In all cases, ****$p < 0.0001$.

### Supplementary Figure S3

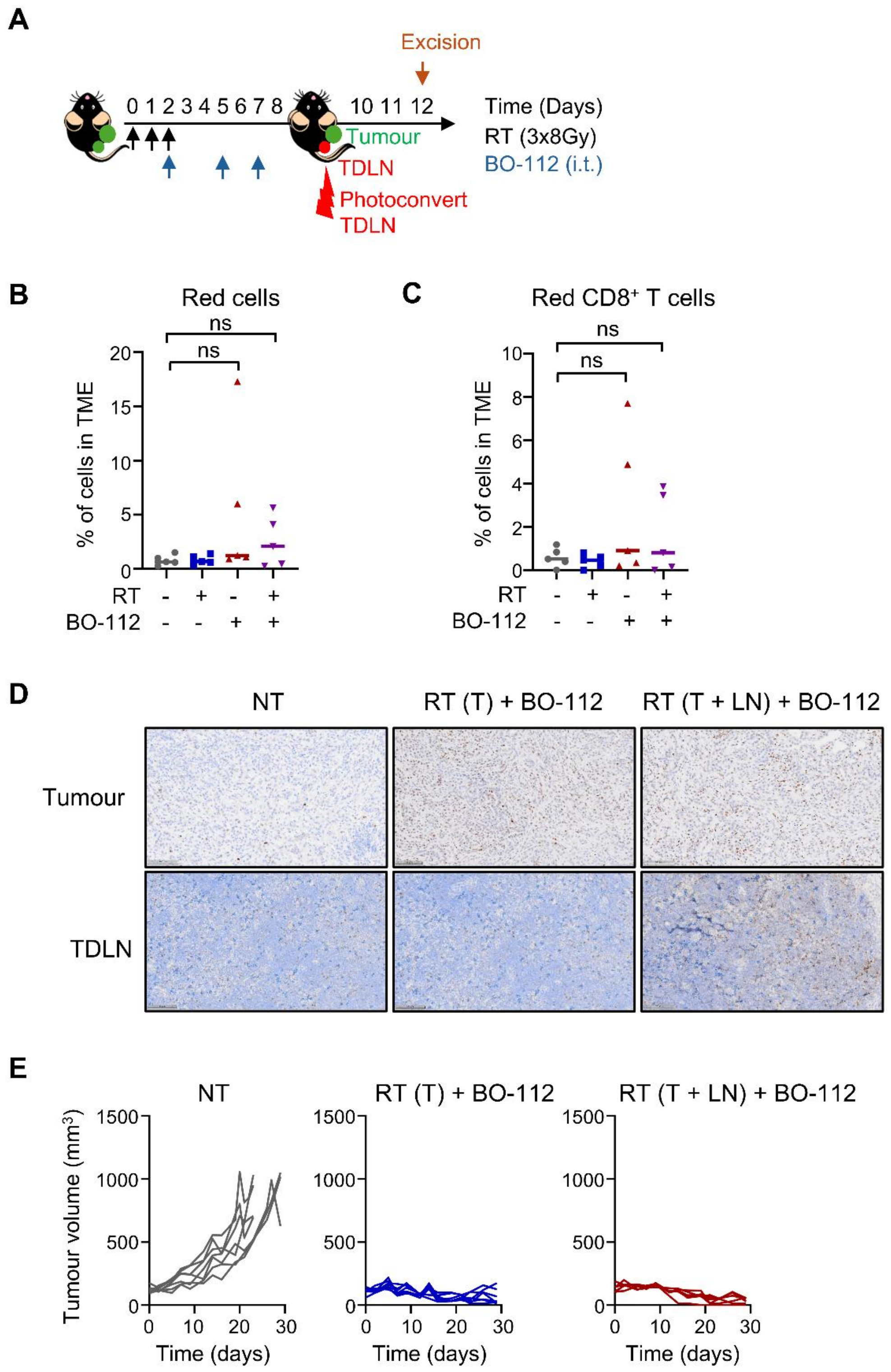

**Supplementary Figure S3.** Additional evidence supporting a dominant role for tumour in situ $CD8^+$ T cells in RT–BO-112 therapeutic efficacy. Related to Figure 2.
**A,** Experimental schematic illustrating that KikGR photoconvertible mice were inoculated with KP475 NSCLC cells subcutaneously and treated with RT–BO-112 dual therapy. Arrows indicate time relative to the first RT fraction. On day 9 following the initiation of RT, tumour-draining lymph nodes (TDLNs) were surgically exposed and photoconverted using violet light. Tumours were excised on day 12 for downstream analysis. **B,** Frequency of photoconverted (red) cells and **C,** frequency of photoconverted $CD8^+$ T cells specifically from the tumours, analysed using flow cytometry. Data shown as mean ± SEM. Each dot represents an individual mouse (n = 5 per group). **D,** Representative IHC images of tumours and TDLNs stained for γH2AX (brown) with nuclear haematoxylin counterstaining (blue), following treatment of KP475 tumour-bearing mice with BO-112 and either local RT targeting the tumour only [RT (T)] or both the tumour and the TDLN [RT (T + LN)]. **E,** Individual tumour growth curves for KP475 tumour-bearing mice receiving no treatment (NT, n = 7), or BO-112 plus RT targeting the tumour only [RT (T) + BO-112, n = 7] or RT targeting both the tumour and TDLN [RT (T + LN) + BO-112, n = 6]. Tumour volumes ranged from 60–188 $mm^3$ at the initiation of RT. Time refers to days with respect to RT initiation. Statistical analysis of flow cytometry data was performed using one-way ANOVA with Dunnett's multiple comparisons test. In all cases, ns = not significant.

**Supplementary Figure S4**

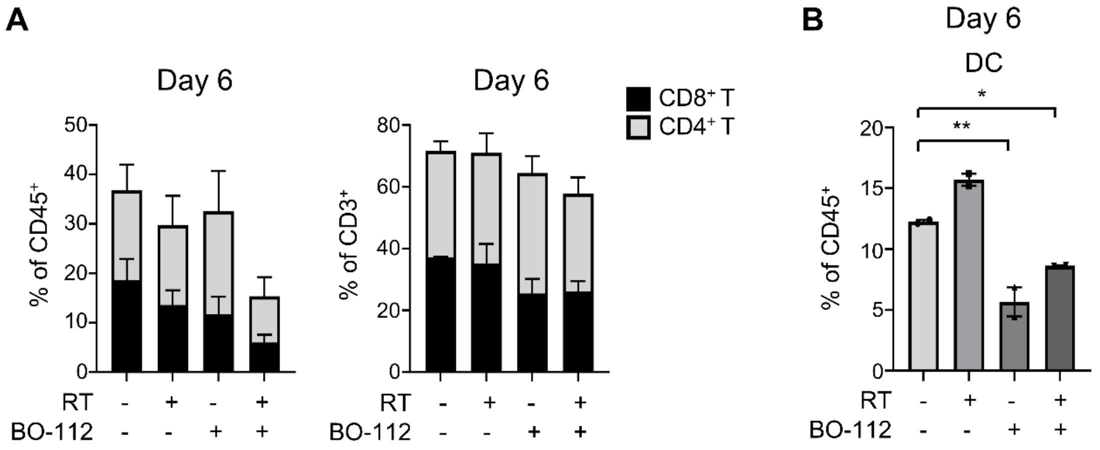


**Supplementary Figure S4.** RT and BO-112 dual therapy alters the early intratumoural immune composition. Related to Figure 3.

Mice bearing subcutaneous KP475 NSCLC tumours were treated with 5% dextrose vehicle (RT–, BO-112–), RT + 5% dextrose (RT+, BO-112–), BO-112 (RT–, BO-112+) or RT + BO-112 dual therapy (RT+, BO-112+). Tumours were surgically harvested for downstream analysis on day 6 following the initiation of treatment. **A,** Frequency of $CD8^+$ and $CD4^+$ T cells within the $CD45^+$ immune compartment (left) and $CD3^+$ lymphocytes (right) (mean ± SEM; n = 3–4). **B,** Frequency of DC within $CD45^+$ immune cells (mean ± SEM; n = 2).

**Supplementary Figure S5**

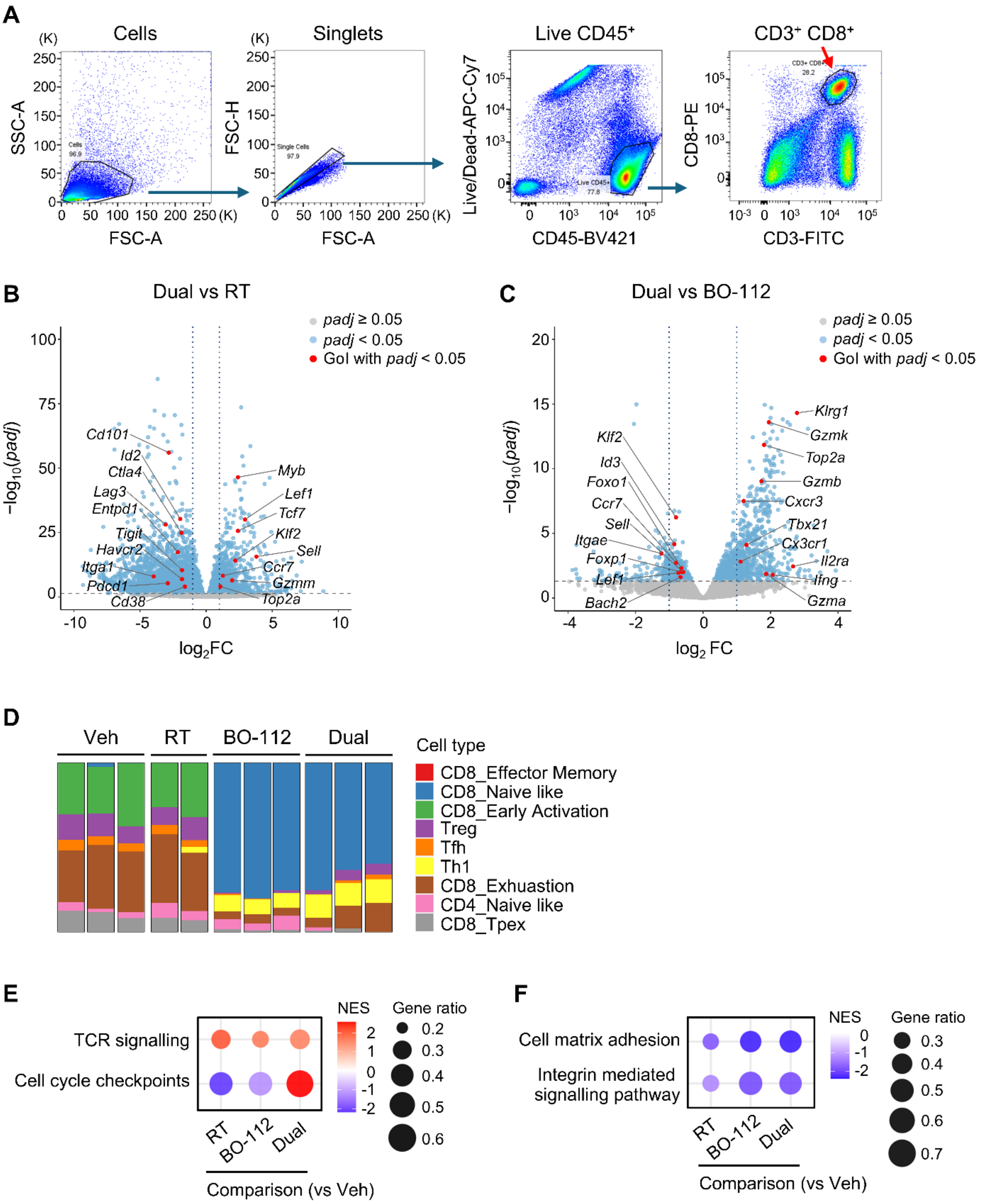

**Supplementary Figure S5.** Transcriptomic and pathway analyses of intratumoural $CD8^+$ T cells. Related to Figure 4.

Mice were implanted with KP475 NSCLC tumours and treated with 5% dextrose vehicle (Veh), RT + Veh (RT), BO-112 or RT + BO-112 (Dual). Live singlet $CD8^+$ T cells were isolated by FACS from tumours on day 12 following the first RT fraction, and bulk RNA was extracted for sequencing. Each sample was pooled from 3–7 tumours. Data shown represent n = 2–3 samples per group. **A,** Representative FACS gating strategy for isolating live singlet $CD45^+$ $CD3^+$ $CD8^+$ T cells for bulk RNA sequencing. Single-cell suspensions were prepared from enzymatically digested KP475 tumours and stained with Near-IR (APC-Cy7) viability dye, CD45-BV421, CD3-FITC and CD8-PE. Cells were sequentially gated on FSC/SSC, singlets (FSC-A/FSC-H), live $CD45^+$ events, and $CD3^+$ and $CD8^+$ co-expression. Volcano plots showing differential gene expression in $CD8^+$ T cells comparing **B,** Dual vs RT and **C,** Dual vs BO-112. The x-axis shows $\log_2$ fold change ($\log_2$FC), and the y-axis shows $-\log_{10}$ of the adjusted *p* value (*padj*). Grey dots: genes with $padj \geq 0.05$; blue dots: genes with $padj < 0.05$; red dots: selected genes of interest (GOI) with $padj < 0.05$. **D,** CIBERSORTx-based deconvolution of bulk RNA-seq data using a reference signature matrix derived from the ProjecTILs murine single-cell RNA-seq atlas of tumour-infiltrating T cells. Stacked bar plots show the inferred proportions of $CD8^+$ T cell subsets, including effector memory, naive-like, early activation, progenitor exhausted (Tpex), and exhausted cells, as well as transcriptionally distinct populations such as Tregs, Th1, Tfh, and $CD4^+$-like subsets. Each bar represents one pooled sample. Detection of $CD4^+$-associated signatures likely reflects transcriptomic heterogeneity within intratumoural $CD8^+$ T cells. **E**–**F,** Bubble plots of GSEA on bulk RNA-seq data using **E,** MSigDB C2: Reactome and **F,** MSigDB C5: Gene Ontology:Biological Processes (GO:BP) collections. Pathways include TCR signalling, cell cycle checkpoints, cell matrix adhesion and integrin-mediated signalling (immune positioning/structural interaction). NES ranges from –2 (blue) to 2 (red); bubble size represents gene ratio (0.2–0.7). Only pathways with $padj < 0.25$ are shown.

**Supplementary Figure S6**

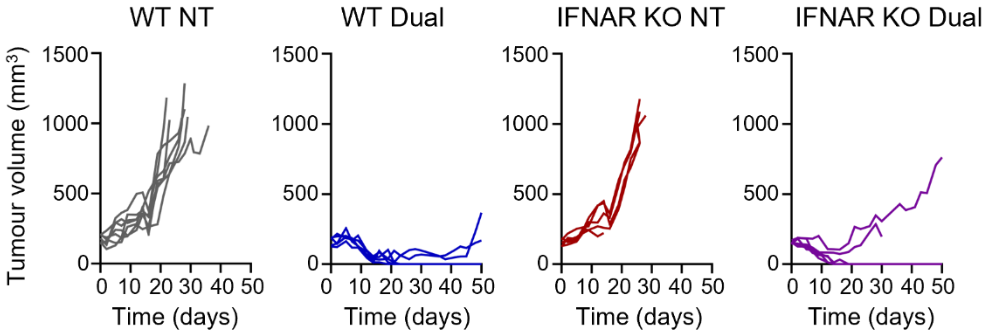


**Supplementary Figure S6.** RT–BO-112 dual therapy efficacy is independent of host type I IFN signalling. Related to Figure 5.

Wildtype (WT) and IFNAR knockout (IFNAR KO) C57BL/6 mice were implanted with KP475 tumours and received no therapy (NT; WT n = 7, KO n = 5) or RT + BO-112 (Dual; WT n = 5, KO n = 6). Individual tumour growth curves are shown. Tumour volumes ranged from 137–212 $mm^3$ at RT initiation. Time on the x-axis indicates days following the first RT fraction.

**Supplementary Figure S7**

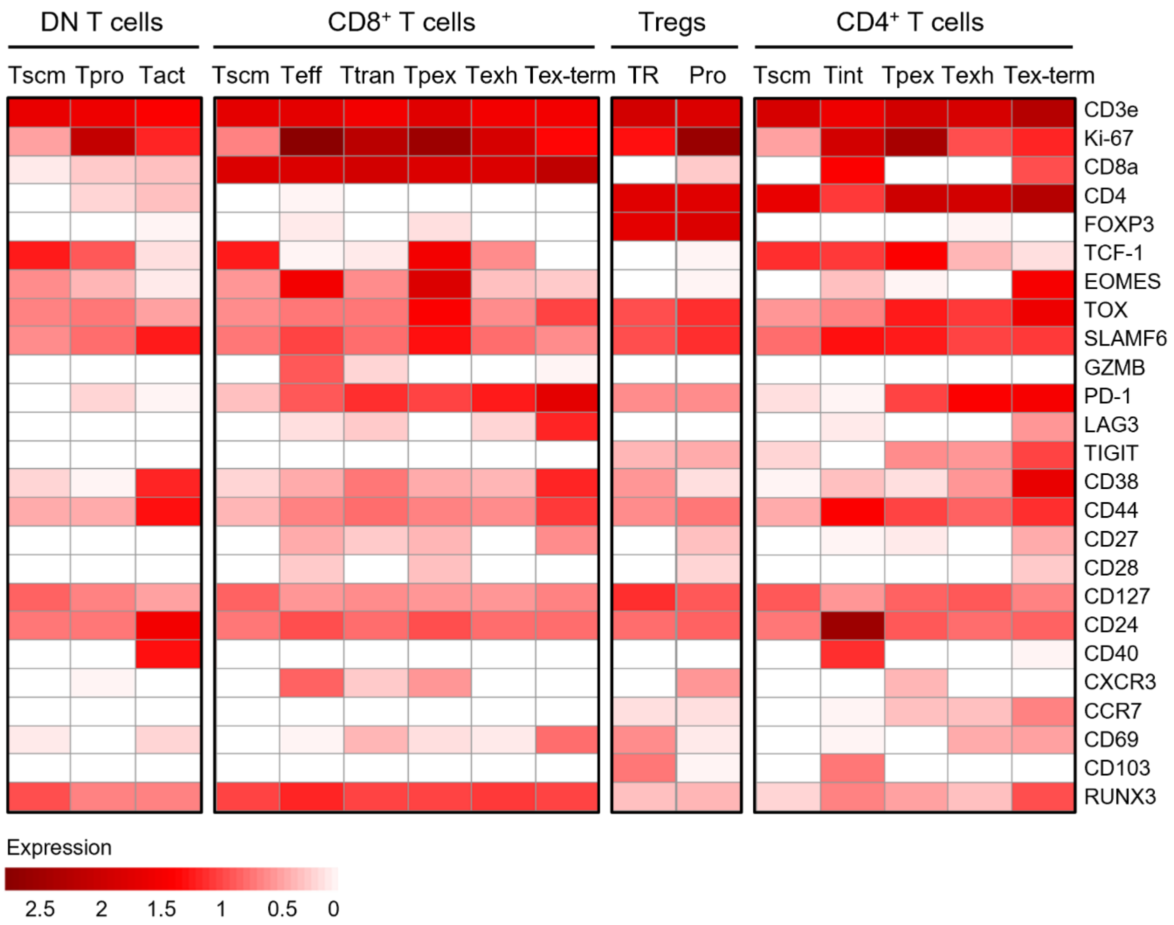


**Supplementary Figure S7.** Marker expression across T-cell clusters identified by mass cytometry (CyTOF). Related to Figure 6.

Mice were implanted with KP475 NSCLC tumours and treated with vehicle (Veh), RT + Veh (RT), BO-112 or RT + BO-112 (Dual). Tumours were harvested on day 12 following the first dose of RT, enzymatically dissociated into single-cell suspensions and pooled per condition (4–5 tumours per group). A total of $3 \times 10^6$ cells per group were stained with a panel of metal-conjugated antibodies and analysed using CyTOF. Heatmap shows the median expression of 25 immune markers across the T cell clusters defined in Figure 6B. $CD3^+$ T cells from KP475 tumours were clustered using FlowSOM and annotated into subsets of double-negative (DN), $CD8^+$, Regulatory (Treg), and $CD4^+$ T cells. Subtypes include stem-like memory (Tscm), proliferative (Tpro), activated (Tact), effector (Teff), transitory (Ttran), progenitor exhausted (Tpex), exhausted (Texh), terminally exhausted (Tex-term), tissue-resident (TR), and intermediate (Tint) populations. Marker intensities are scaled across clusters, with higher expression levels represented by deeper red colour intensity.

Supplementary Tables

| Time point | Compartment | Cell type | RT vs Veh | BO-112 vs Veh | Dual vs Veh |
|---|---|---|---|---|---|
| Day 6 | CD45⁺ | CD8⁺ T | 0.543 | 0.326 | 0.034 |
| | | CD4⁺ T | 0.991 | 0.976 | 0.563 |
| | | DC | 0.045 | 0.005 | 0.039 |
| Day 12 | CD45⁺ | CD8⁺ T | 0.155 | 0.108 | 0.010 |
| | | CD4⁺ T | 0.817 | 0.352 | 0.023 |
| | | DC | 0.107 | 0.0002 | < 0.0001 |
| Day 12 | CD3⁺ | CD8⁺ T | 0.023 | 0.013 | 0.0003 |
| | | CD4⁺ T | 0.756 | 0.219 | 0.003 |

**Supplementary Table S1**. *P*-values using one-way ANOVA with Dunnett's or Tukey's multiple comparisons tests for Figure 3A and B, and Supplementary Figure S4A.

| | |
|---|---|
| Gene symbol | *SELL, LEF1, TCF7, KLF2, CCR7, ID3, BACH2, IL7R, FOXO1, IKZF1, CXCR5, SLAMF6, BTLA, CD27* |

**Supplementary Table S2**. Tscm gene signature derived from preclinical RNA sequencing of intratumoural $CD8^+$ T cells. Related to Figure 7.

| Figure & panel | Target marker | Antibody clone | Fluorochrome | Supplier | Catalogue# |
|---|---|---|---|---|---|
| Fig. 3a-c, f, g | CD45 | 30F11 | BV421 | BioLegend | 103134 |
| | CD3ε | 145-2C11 | PerCP-Cy5.5 | BioLegend | 100328 |
| | CD4 | RM4-5 | BV785 | BioLegend | 100552 |
| | CD25 | PC61 | BV711 | BioLegend | 102049 |
| | FOXP3 | FJK-16s | PE | Invitrogen | 12-5773-82 |
| | CD8α | 53-6.7 | Alexa Fluor 700 | BioLegend | 100730 |
| | MHC II | M5/114.15.2 | BV510 | BioLegend | 107636 |
| | CD11c | N418 | PE-Cy7 | Invitrogen | 25-0114-82 |
| Fig. 3d | CD45 | 30F11 | BV785 | BioLegend | 103149 |
| | CD3ε | 17A2 | BUV395 | BD Bioscience | 563565 |
| | CD8α | 53-6.7 | Alexa Fluor 700 | BioLegend | 100730 |
| | Granzyme B | QA16A02 | FITC | BioLegend | 372206 |

| | | | | | |
|---|---|---|---|---|---|
| Supplementary Fig. 3b, c | CD45 | 30F11 | BV 510 | BioLegend | 103138 |
| | CD3ε | 17A2 | BUV395 | BD Bioscience | 563565 |
| | CD8α | 53-6.7 | Alexa Fluor 700 | BioLegend | 100730 |
| Supplementary Fig. 4a, b | CD45 | 30F11 | BV421 | BioLegend | 103134 |
| | CD3ε | 145-2C11 | FITC | BioLegend | 100306 |
| | CD4 | RM4-5 | BV785 | BioLegend | 100552 |
| | CD8α | 53-6.7 | PE | BioLegend | 100708 |
| | MHC II | M5/114.15.2 | BV510 | BioLegend | 107636 |
| | CD11c | N418 | APC | BioLegend | 117310 |
| Supplementary Fig. 5a | CD45 | 30F11 | BV421 | BioLegend | 103134 |
| | CD3ε | 145-2C11 | FITC | BioLegend | 100306 |
| | CD8α | 53-6.7 | PE | BioLegend | 100708 |

**Supplementary Table S3**. Flow cytometry panels and antibodies.

| | BO-112 vs Veh | | Dual vs Veh | | RT vs Veh | | Dual vs BO-112 | | Dual vs RT | |
|---|---|---|---|---|---|---|---|---|---|---|
| gene | $log_2FC$ | padj | $log_2FC$ | padj | $log_2FC$ | padj | $log_2FC$ | padj | $log_2FC$ | padj |
| *Apc* | 0.60 | 0.04 | 0.33 | 0.32 | 0.44 | 0.17 | -0.22 | 0.61 | -0.09 | 0.85 |
| *Aurkb* | -1.06 | 0.00 | 0.97 | 0.00 | -0.82 | 0.00 | 2.02 | 0.00 | 1.73 | 0.00 |
| *Axin2* | 1.42 | 0.00 | 0.62 | 0.05 | 0.28 | 0.58 | -0.74 | 0.06 | 0.35 | 0.42 |
| *Bach2* | 1.60 | 0.00 | 0.93 | 0.00 | 0.46 | 0.04 | -0.67 | 0.02 | 0.52 | 0.09 |
| *Bcl9* | 1.67 | 0.00 | 1.13 | 0.00 | -0.32 | 0.69 | -0.51 | 0.29 | 1.50 | 0.00 |
| *Btla* | 2.31 | 0.00 | 2.19 | 0.00 | -0.26 | 0.73 | -0.08 | 0.84 | 2.46 | 0.00 |
| *Ccnb1* | -0.88 | 0.01 | 1.37 | 0.00 | -0.73 | 0.00 | 2.22 | 0.00 | 2.06 | 0.00 |
| *Ccne1* | -0.39 | 0.37 | 1.22 | 0.00 | -0.38 | 0.39 | 1.63 | 0.00 | 1.60 | 0.00 |
| *Ccr7* | 1.79 | 0.00 | 0.96 | 0.00 | -0.27 | 0.03 | -0.80 | 0.00 | 1.27 | 0.00 |
| *Cd101* | -2.64 | 0.00 | -2.83 | 0.00 | -0.02 | 0.95 | -0.15 | 0.68 | -2.83 | 0.00 |
| *Cd160* | -1.91 | 0.00 | -0.89 | 0.00 | -0.22 | 0.25 | 1.07 | 0.00 | -0.64 | 0.03 |
| *Cd24a* | -0.96 | 0.00 | -1.38 | 0.00 | -0.41 | 0.04 | -0.38 | 0.28 | -0.94 | 0.00 |
| *Cd27* | 0.96 | 0.00 | 1.03 | 0.00 | 0.18 | 0.42 | 0.11 | 0.64 | 0.87 | 0.00 |
| *Cd38* | -2.40 | 0.00 | -1.48 | 0.00 | 0.16 | 0.71 | 1.00 | 0.07 | -1.61 | 0.00 |
| *Cd44* | -2.49 | 0.00 | -1.77 | 0.00 | 0.55 | 0.00 | 0.76 | 0.00 | -2.30 | 0.00 |
| *Cenpa* | -0.84 | 0.00 | 0.18 | 0.48 | 0.00 | 1.00 | 1.05 | 0.00 | 0.19 | 0.54 |
| *Ctla4* | -1.97 | 0.00 | -1.27 | 0.00 | 0.61 | 0.00 | 0.73 | 0.00 | -1.87 | 0.00 |

| | | | | | | | | | | |
|---|---|---|---|---|---|---|---|---|---|---|
| *Ctnnbip1* | -1.87 | 0.00 | -1.13 | 0.00 | -0.28 | 0.60 | 0.76 | 0.18 | -0.85 | 0.04 |
| *Cx3cr1* | -4.69 | 0.00 | -3.62 | 0.00 | 0.29 | 0.17 | 1.11 | 0.00 | -3.89 | 0.00 |
| *Cxcl10* | 1.44 | 0.00 | 1.72 | 0.00 | 0.82 | 0.00 | 0.32 | 0.21 | 0.92 | 0.00 |
| *Cxcr3* | -0.51 | 0.02 | 0.68 | 0.00 | 0.01 | 0.99 | 1.20 | 0.00 | 0.62 | 0.02 |
| *Cxcr4* | 1.23 | 0.00 | 0.95 | 0.00 | 0.06 | 0.85 | -0.24 | 0.20 | 0.89 | 0.00 |
| *Cxcr5* | 2.51 | 0.00 | 2.90 | 0.00 | -0.78 | 0.16 | 0.38 | 0.37 | 3.63 | 0.00 |
| *Dhx58* | 2.63 | 0.00 | 2.29 | 0.00 | 0.31 | 0.49 | -0.30 | 0.25 | 2.05 | 0.00 |
| *Dkk2* | -5.69 | 0.00 | -6.91 | 0.00 | 0.52 | 0.76 | -1.18 | NA | -7.40 | 0.00 |
| *Egr2* | -0.56 | 0.01 | -1.34 | 0.00 | 0.02 | 0.98 | -0.74 | 0.02 | -1.31 | 0.00 |
| *Egr3* | -0.75 | 0.00 | -1.35 | 0.00 | -0.09 | 0.83 | -0.54 | 0.03 | -1.15 | 0.00 |
| *Entpd1* | -2.85 | 0.00 | -1.77 | 0.00 | 0.38 | 0.10 | 1.10 | 0.00 | -2.14 | 0.00 |
| *Eomes* | 1.20 | 0.00 | 1.90 | 0.00 | -0.07 | 0.89 | 0.75 | 0.01 | 2.00 | 0.00 |
| *Foxo1* | 1.25 | 0.00 | 0.59 | 0.01 | 0.73 | 0.00 | -0.64 | 0.00 | -0.10 | 0.77 |
| *Foxo3* | 1.22 | 0.00 | 0.38 | 0.15 | 0.22 | 0.45 | -0.81 | 0.01 | 0.21 | 0.56 |
| *Foxp1* | 1.86 | 0.00 | 1.17 | 0.00 | 0.09 | 0.82 | -0.66 | 0.01 | 1.14 | 0.00 |
| *Gzma* | -1.07 | 0.04 | 0.75 | 0.19 | 0.20 | 0.52 | 1.87 | 0.01 | 0.60 | 0.41 |
| *Gzmb* | -0.74 | 0.00 | 0.97 | 0.00 | 0.56 | 0.00 | 1.73 | 0.00 | 0.42 | 0.20 |
| *Gzmk* | 0.30 | 0.30 | 2.24 | 0.00 | 0.19 | 0.62 | 1.95 | 0.00 | 2.06 | 0.00 |
| *Gzmm* | 1.66 | 0.00 | 1.93 | 0.00 | 0.07 | 0.95 | 0.35 | 0.48 | 1.97 | 0.00 |
| *Havcr2* | -3.02 | 0.00 | -0.89 | 0.01 | 0.94 | 0.00 | 2.12 | 0.00 | -1.83 | 0.00 |
| *Icos* | -1.09 | 0.00 | -0.71 | 0.00 | 0.74 | 0.00 | 0.42 | 0.01 | -1.45 | 0.00 |
| *Id2* | -1.72 | 0.00 | -1.22 | 0.00 | 0.78 | 0.00 | 0.55 | 0.01 | -1.96 | 0.00 |
| *Id3* | 1.61 | 0.00 | 0.72 | 0.00 | -0.11 | 0.88 | -0.85 | 0.00 | 0.82 | 0.00 |
| *Ifih1* | 1.89 | 0.00 | 1.76 | 0.00 | 0.16 | 0.74 | -0.09 | 0.78 | 1.64 | 0.00 |
| *Ifit1* | 3.26 | 0.00 | 2.70 | 0.00 | 0.65 | 0.06 | -0.53 | 0.05 | 2.09 | 0.00 |
| *Ifit3* | 3.31 | 0.00 | 2.76 | 0.00 | 0.67 | 0.06 | -0.51 | 0.03 | 2.17 | 0.00 |
| *Ifnar2* | 0.93 | 0.00 | 0.65 | 0.00 | -0.08 | 0.92 | -0.24 | 0.45 | 0.78 | 0.00 |
| *Ifng* | -2.85 | 0.00 | -0.76 | 0.22 | 0.73 | 0.00 | 2.07 | 0.02 | -1.50 | 0.02 |
| *Ikzf1* | 1.42 | 0.00 | 1.21 | 0.00 | 0.11 | 0.57 | -0.16 | 0.43 | 1.12 | 0.00 |
| *Il2ra* | -3.42 | 0.00 | -0.55 | 0.55 | 0.68 | 0.00 | 2.67 | 0.00 | -1.36 | 0.12 |
| *Il7r* | 1.37 | 0.00 | 0.48 | 0.07 | 0.56 | 0.00 | -0.85 | 0.01 | -0.06 | 0.90 |
| *Irf7* | 3.86 | 0.00 | 3.17 | 0.00 | 0.50 | 0.02 | -0.66 | 0.01 | 2.71 | 0.00 |
| *Irf9* | 2.07 | 0.00 | 1.55 | 0.00 | 0.32 | 0.20 | -0.49 | 0.02 | 1.27 | 0.00 |
| *Isg15* | 2.74 | 0.00 | 2.35 | 0.00 | 0.65 | 0.02 | -0.36 | 0.18 | 1.79 | 0.00 |
| *Itga1* | -4.60 | 0.00 | -3.57 | 0.00 | 0.35 | 0.02 | 0.97 | 0.18 | -3.97 | 0.00 |

| | | | | | | | | | |
|---|---|---|---|---|---|---|---|---|---|
| *Itga4* | 1.64 | 0.00 | 1.91 | 0.00 | 0.12 | 0.93 | 0.31 | 0.12 | 1.82 | 0.00 |
| *Itgae* | -0.15 | 0.71 | -1.48 | 0.00 | -0.29 | 0.23 | -1.23 | 0.00 | -1.24 | 0.00 |
| *Itgb1* | -2.28 | 0.00 | -1.82 | 0.00 | -0.01 | 0.99 | 0.48 | 0.05 | -1.84 | 0.00 |
| *Jak1* | 0.99 | 0.00 | 0.65 | 0.00 | 0.22 | 0.08 | -0.29 | 0.17 | 0.46 | 0.00 |
| *Klf2* | 2.86 | 0.00 | 2.01 | 0.00 | -0.33 | 0.76 | -0.80 | 0.00 | 2.22 | 0.00 |
| *Klf4* | 0.97 | 0.00 | 0.28 | 0.43 | 0.25 | 0.28 | -0.61 | 0.10 | 0.01 | 0.99 |
| *Klf7* | 1.17 | 0.00 | 0.63 | 0.00 | 0.14 | 0.78 | -0.50 | 0.08 | 0.54 | 0.04 |
| *Klrg1* | 0.15 | 0.89 | 3.10 | 0.00 | 0.73 | 0.09 | 2.78 | 0.00 | 2.15 | 0.03 |
| *Lag3* | -3.56 | 0.00 | -2.58 | 0.00 | 0.50 | 0.00 | 0.99 | 0.00 | -3.07 | 0.00 |
| *Lef1* | 3.97 | 0.00 | 3.16 | 0.00 | 0.20 | 0.59 | -0.73 | 0.01 | 2.94 | 0.00 |
| *Lig1* | -1.35 | 0.00 | 0.06 | 0.86 | -0.41 | 0.00 | 1.42 | 0.00 | 0.45 | 0.10 |
| *Ly6a* | 0.71 | 0.00 | 1.37 | 0.00 | 0.46 | 0.00 | 0.70 | 0.00 | 0.93 | 0.00 |
| *Mki67* | -1.61 | 0.00 | 0.86 | 0.02 | -0.10 | 0.97 | 2.51 | 0.00 | 0.79 | 0.07 |
| *Mx1* | 2.06 | 0.00 | 1.86 | 0.00 | 0.99 | 0.04 | -0.16 | 0.76 | 0.98 | 0.01 |
| *Myb* | 2.69 | 0.00 | 2.49 | 0.00 | 0.05 | 0.95 | -0.16 | 0.42 | 2.41 | 0.00 |
| *Nfat5* | -1.03 | 0.00 | -1.17 | 0.00 | 0.29 | 0.08 | -0.09 | 0.75 | -1.43 | 0.00 |
| *Oas2* | 4.63 | 0.00 | 4.17 | 0.00 | 0.67 | 0.67 | -0.44 | 0.20 | 3.62 | 0.00 |
| *Pdcd1* | -4.44 | 0.00 | -2.66 | 0.00 | 0.22 | 0.27 | 1.67 | 0.04 | -2.90 | 0.00 |
| *Plk1* | -0.57 | 0.11 | 1.51 | 0.00 | -0.77 | 0.00 | 2.09 | 0.00 | 2.26 | 0.00 |
| *Ppargc1b* | 1.82 | 0.00 | 1.17 | 0.00 | 0.00 | 1.00 | -0.62 | 0.17 | 1.21 | 0.00 |
| *Prf1* | -1.28 | 0.00 | -0.51 | 0.03 | 0.73 | 0.12 | 0.81 | 0.00 | -1.21 | 0.00 |
| *Prkaa1* | 0.85 | 0.00 | 0.63 | 0.00 | 0.10 | 0.81 | -0.18 | 0.43 | 0.54 | 0.00 |
| *Ptk2* | -1.19 | 0.00 | -1.98 | 0.00 | -0.67 | 0.06 | -0.76 | 0.16 | -1.26 | 0.00 |
| *Rasgrp2* | 3.33 | 0.00 | 3.33 | 0.00 | 0.41 | 0.37 | 0.04 | 0.88 | 2.86 | 0.00 |
| *Rictor* | 1.15 | 0.00 | 0.70 | 0.00 | 0.30 | 0.21 | -0.40 | 0.19 | 0.42 | 0.11 |
| *Rigi* | 2.25 | 0.00 | 1.87 | 0.00 | 0.28 | 0.30 | -0.34 | 0.10 | 1.61 | 0.00 |
| *Rrm2* | -0.90 | 0.00 | 1.12 | 0.00 | -0.53 | 0.00 | 2.03 | 0.00 | 1.64 | 0.00 |
| *Rsad2* | 3.67 | 0.00 | 3.05 | 0.00 | 0.76 | 0.10 | -0.59 | 0.03 | 2.33 | 0.00 |
| *S1pr1* | 3.34 | 0.00 | 2.45 | 0.00 | 0.11 | 0.78 | -0.81 | 0.00 | 2.29 | 0.00 |
| *Sell* | 4.21 | 0.00 | 3.53 | 0.00 | -0.48 | 0.69 | -0.59 | 0.01 | 3.80 | 0.00 |
| *Sfrp1* | -3.00 | 0.00 | -3.24 | 0.00 | -1.74 | 0.14 | -0.17 | NA | -1.58 | 0.27 |
| *Sfrp2* | -3.62 | 0.00 | -4.44 | 0.00 | -0.69 | 0.69 | -0.77 | NA | -3.59 | 0.00 |
| *Sirt1* | 1.39 | 0.00 | 0.80 | 0.00 | 0.55 | 0.00 | -0.56 | 0.04 | 0.27 | 0.28 |
| *Slamf6* | 2.43 | 0.00 | 2.73 | 0.00 | 0.09 | 0.84 | 0.34 | 0.13 | 2.65 | 0.00 |
| *Sox4* | -0.71 | 0.01 | -1.74 | 0.00 | -0.24 | 0.34 | -1.03 | 0.00 | -1.40 | 0.00 |

| | | | | | | | | | |
|---|---|---|---|---|---|---|---|---|---|
| *Stat1* | 1.64 | 0.00 | 1.53 | 0.00 | 0.12 | 0.74 | -0.07 | 0.81 | 1.44 | 0.00 |
| *Stat2* | 1.47 | 0.00 | 1.16 | 0.00 | -0.14 | 0.82 | -0.28 | 0.32 | 1.38 | 0.00 |
| *Stat3* | -1.10 | 0.00 | -1.08 | 0.00 | 0.49 | 0.00 | 0.06 | 0.85 | -1.55 | 0.00 |
| *Tbk1* | 0.76 | 0.00 | 0.77 | 0.00 | 0.42 | 0.09 | 0.05 | 0.85 | 0.37 | 0.03 |
| *Tbx21* | -2.23 | 0.00 | -0.98 | 0.00 | 0.24 | 0.15 | 1.29 | 0.00 | -1.20 | 0.00 |
| *Tcf7* | 2.87 | 0.00 | 2.30 | 0.00 | -0.13 | 0.70 | -0.51 | 0.04 | 2.40 | 0.00 |
| *Tigit* | -2.45 | 0.00 | -1.38 | 0.00 | 0.45 | 0.01 | 1.07 | 0.00 | -1.82 | 0.00 |
| *Tle1* | 0.90 | 0.00 | 0.12 | 0.75 | -0.03 | 0.98 | -0.74 | 0.05 | 0.18 | 0.67 |
| *Tle4* | 1.78 | 0.00 | 1.27 | 0.00 | 0.18 | 0.63 | -0.47 | 0.08 | 1.12 | 0.00 |
| *Tnfrsf4* | -3.89 | 0.00 | -2.43 | 0.00 | 0.20 | 0.35 | 1.40 | 0.01 | -2.64 | 0.00 |
| *Tnfrsf9* | -3.58 | 0.00 | -2.27 | 0.00 | 0.72 | 0.00 | 1.30 | 0.06 | -2.98 | 0.00 |
| *Top2a* | -1.17 | 0.00 | 0.64 | 0.01 | -0.48 | 0.00 | 1.81 | 0.00 | 1.09 | 0.00 |
| *Traf3* | 0.52 | 0.00 | 0.25 | 0.12 | 0.31 | 0.11 | -0.23 | 0.36 | -0.05 | 0.86 |
| *Vcam1* | -3.86 | 0.00 | -4.72 | 0.00 | -0.81 | 0.00 | -0.82 | 0.35 | -3.89 | 0.00 |
| *Zeb1* | 1.06 | 0.00 | 0.66 | 0.00 | -0.02 | 0.98 | -0.37 | 0.24 | 0.71 | 0.00 |
| *Zeb2* | -2.68 | 0.00 | -1.94 | 0.00 | 0.53 | 0.02 | 0.83 | 0.13 | -2.42 | 0.00 |
| *Znrf3* | 1.67 | 0.00 | 0.79 | 0.01 | 0.00 | 1.00 | -0.82 | 0.00 | 0.78 | 0.03 |

**Supplementary Table S4.** $Log_2$ fold change ($log_2FC$) and adjusted *p* values (*padj*) for genes of interest across five comparisons from bulk RNAseq of intratumoural $CD8^+$ T cells. "Dual" indicates combined therapy with Radiotherapy (RT; 3 x 8Gy) and BO-112, whereas "RT" indicates 3 x 8Gy delivered with vehicle control.

| Target marker | Antibody clone | Metal conjugate | Supplier | Catalogue# |
|---|---|---|---|---|
| CD3e | 17A2 | 173Yb | Biolegend | 101202 |
| Ki67 | S0A15 | 152Sm | ThermoFisher | 14-5698-82 |
| CD8a | 53-6.7 | 153Eu | Standard Biotools | 3153012B |
| CD4 | RM4-5 | 145Nd | Standard Biotools | 3145002B |
| FoxP3 | FJK-16s | 158Gd | Standard Biotools | 3158003A |
| TCF-1 | C63D9 | 147Sm | Cell Signaling Technology | 2203 |
| EOMES | Dan11mag | 156Gd | ThermoFisher | 14-4875-82 |
| TOX | NAN448B | 166Er | BD Bioscience | 569634 |
| SLAMF6 | TC15-12F12.2 | 195Pt | Biolegend | 115902 |
| Granzyme B | QA16A02 | 141Pr | Biolegend | 396402 |
| PD-1 | 29F.1A12 | 159Tb | Standard Biotools | 3159024B |

| | | | | |
|---|---|---|---|---|
| LAG3 | C9B7W | 151Eu | Biolegend | 125202 |
| TIGIT | 3G9 | 146Nd | Biolegend | 142102 |
| CD38 | 90 | 175Lu | Standard Biotools | 3175014B |
| CD44 | 1M7 | 144Nd | Biolegend | 103002 |
| CD27 | LG.3A10 | 150Nd | Standard Biotools | 3150017B |
| CD28 | 37.51 | 157Gd | Biolegend | 102119 |
| CD127 | A7R34 | 174Yb | Standard Biotools | 3174013B |
| CD24 | M1/69 | 194Pt | Biolegend | 101802 |
| CD40 | HM440 | 176Yb | Biolegend | 102902 |
| CXCR3 | CXCR3-173 | 139La | Biolegend | 126502 |
| CCR7 | 4B12 | 161Dy | Biolegend | 120101 |
| CD69 | H1-2F3 | 143Nd | Standard Biotools | 3143004B |
| CD103 | 2E7 | 164Dy | Biolegend | 121402 |
| RUNX3 | R3-5G4 | 170Er | BD Bioscience | 564813 |

**Supplementary Table S5**. CyTOF antibodies.

Supplementary Methods and Materials

**Cell lines**

KP475 (*Kras*$^{G12D}$, *Trp53*$^{-/-}$) murine non-small cell lung cancer (NSCLC) cells (a gift from Dr Carla Martins, CRUK Cambridge Centre and University of Cambridge) were cultured in F-12/DMEM medium (Gibco), and MB49 murine bladder cancer cells (a gift from Prof. Michael O'Donnell, University of Iowa, Iowa City, IA, USA) were cultured in DMEM (Gibco), both supplemented with 10% foetal bovine serum (FBS; Gibco). Cells were maintained at 37 °C in a humidified incubator with 5% $CO_2$. Both cell lines were routinely tested for mycoplasma contamination and authenticated based on morphology and growth characteristics.

**Mouse models**

C57BL/6 wild-type mice were purchased from Envigo (females, 6–8 weeks old, for the KP475 NSCLC model; males, 3–4 weeks old, for the MB49 bladder cancer model). Kikume Green:Red photoconvertible mice ((CAG-KikGR)33Hadj/J) were purchased from Jax. Both KikGR and IFNAR KO mice were bred in-house at the CRUK Manchester Institute (CRUK MI), University of Manchester. Knockout status was confirmed by genotyping. Both male and female transgenic mice were used. Mice were given at least one week to acclimatise to the facility. Mice were housed at a maximum density of seven animals per cage in individual ventilated cages under specific pathogen-free conditions. Cages were kept on a 12/12 light/dark cycle and contained aspenchips-2 bedding, sizzlenest nesting material, and a variety of objects for environmental enrichment. Teklad Global 19% protein extruded rodent diet and sterilised water were provided ad libitum. Mice were typically 7-12 weeks of age at the time of tumour implantation. Tumour volume was measured with digital callipers and calculated using the formula: volume = length × width$^2$ × 0.5 and did not exceed 1200 mm$^3$. All animal studies were performed under a United Kingdom Home Office Project license (PCC943F76 and PP1231845) and approved by the CRUK Manchester Institute Animal Welfare and Ethical Review Board (AWERB).

**In vivo treatment**

Subcutaneous tumours were established by injecting 1 × 10$^6$ KP475 or MB49 cells into the right flank of each mouse. Mice were treated when tumour volume reached ~75–200 mm$^3$, typically day 9 post-inoculation for KP475 and day 7 for MB49. Mice (typically 7 per group for a biologically independent experiment) were randomised into groups to ensure comparable tumour volume distribution across cohorts prior to commencing therapy. Radiotherapy (RT) was delivered as three consecutive daily fractions of 8 Gy (3 × 8Gy) using a Pantak HF-320 320 kV X-ray unit (Gulmay Medical), operated at 50 kV, 10 mA, 0.57 mm Al filter; dose rate: 1.15 Gy/min. Mice were shielded in a custom-designed lead jig, leaving only the tumour-bearing flank exposed. In selected experiments, a larger opening was used to simultaneously expose the tumour and the TDLN. BO-112 (48 µg per dose in 80 µl of 0.6 mg/ml; Highlight Therapeutics) or vehicle control (80 µl of 5% dextrose) was administered intratumourally (i.t.) on days 2, 5, and 7, relative to the first RT fraction. Anti-PD-1 monoclonal antibody (clone RMP1-4; Bio X Cell; 200 µg per dose) was administered intraperitoneally (i.p.) on days 0, 2, and 4, relative to first RT fraction. For T cell depletion, anti-CD4 (clone GK1.5; Bio X Cell; 250 µg per dose) or anti-CD8 (clone YTS 169.4; Bio X Cell; 500 µg per dose) antibody was administered intraperitoneally. FTY-720 (Fingolimod; Enzo Life Sciences) was administered daily by oral gavage. An initial dose of 25 µg in 200 µl was followed by daily maintenance doses of 5 µg in 100 µl, continued until experimental endpoint. Vehicle control mice received hydroxypropyl methylcellulose (HPMC; Sigma). Researchers were responsible for group allocation (cohorts for treatment), and all procedures and outcome measurements were conducted by technical staff from the CRUK MI Biological

Resources Unit (BRU), who were unaware of expected outcome. Data analysis was performed by researchers leading the study.

**Photoconversion of tumour draining lymph nodes**

For photoconversion, Kikume Green:Red mice were anaesthetised with isofluorane and draining lymph nodes surgically exposed through a small surgical incision. Exposed lymph nodes were irradiated continuously for 2 min with violet light using a modified 300W compact xenon arc lamp (Excelitas Cermax PE300) fitted with an Edmunds Optics bandpass filter with a centre transmission wavelength of 400 nm at an intensity of 150 $mW/cm^2$. During the violet light emission, the rest of the mouse was covered with black cloth to protect it from photoconversion. Following photoconversion, the incision was closed with glue. Three days post photoconversion mice were humanely killed and tumours, tumour draining lymph nodes and alternative non-draining inguinal lymph nodes harvested for subsequent analysis.

**In vivo tumour rechallenge**

Mice that remained tumour-free for at least 60 days following therapy were rechallenged subcutaneously with tumour cells (same cell number and preparation as in primary implantation) into the contralateral (left) flank. Tumour-naive mice were injected as controls.

**Flow cytometry**

KP475 murine NSCLC tumours were excised, enzymatically dissociated using the Tumour Dissociation Kit (Miltenyi Biotec), and passed through a 70 µm strainer (Miltenyi Biotec) to obtain single-cell suspensions. Cells were stained with a viability dye (Near-IR Dead Cell Stain Kit; 1:1000; Invitrogen) in PBS for 15 min at room temperature, washed with FACS buffer (2% FBS in PBS), blocked with anti-CD16/32 (clone 93; 1:100; eBioscience), and stained with fluorochrome-conjugated extracellular antibodies diluted in Brilliant Staining Buffer (BD Biosciences) at 4 °C for 20 min. Cells were then fixed and permeabilised using True-Nuclear Transcription Factor Buffer Set (BioLegend), re-blocked with anti-CD16/32, and stained for intracellular markers, followed by fixation in 1% paraformaldehyde (BD Cytofix). Antibody panels are listed in Supplementary Table S3.

Single-stain controls using UltraComp eBeads (Invitrogen) or cells, and fluorescence minus one (FMO) controls were included for compensation and gating. Data were acquired on a BD LSRFortessa cytometer (BD Biosciences) and analysed using FlowJo software (BD Biosciences), with ≥ 10,000 live cells acquired per sample.

**Immunohistochemistry (IHC)**

KP475 murine NSCLC tumours were surgically excised, fixed in 10% neutral-buffered formalin, transferred to 70% ethanol, and embedded in paraffin. FFPE sections (4 μm) were stained for CD8a (clone 4SM15; 1:100; eBioscience), Granzyme B (clone EPR22645-206; 1:6000; Abcam) and γH2AX (phospho-Ser139; 1:300; CST, #2577). Multiplex staining for CD8 and Granzyme B was performed using the Opal TSA detection system (Opal 520®, 570®, and 650®; Akoya Biosciences) on the BOND RX automated stainer (Leica Microsystems, Milton Keynes, UK). Slides were counterstained with DAPI (1:1000; #D3571; Thermo Fisher), mounted with ProLong Gold Antifade Mountant (#P36934; Invitrogen), and scanned using the VS-120 slide scanner (Olympus). Chromogenic IHC for γH2AX was performed on the BOND RX using pH 9 EDTA-based antigen retrieval, HRP/DAB detection (Vector Laboratories), and haematoxylin counterstaining, followed by mounting with Pertex (CellPath, #00801). Slides were scanned using the VS-200S scanner (Olympus). Image analysis and quantification were performed using HALO image analysis software with the HighPlex v4.0.5107.4707 module (Indica Labs, Albuquerque, NM, USA).

**Intratumoural $CD8^+$ T cell isolation and RNA extraction**

KP475 murine NSCLC tumours were enzymatically dissociated into single-cell suspensions as described above. $CD8^+$ T cells (Live, $CD45^+$, $CD3^+$, $CD8^+$; Supplementary Table S3) were isolated by fluorescence-activated cell sorting (FACS) using a BD FACSAria III cell sorter. For each sample, tumours from 3–7 mice were pooled, and 3 biological replicates were obtained per condition. Bulk RNA was extracted immediately following sorting using the RNeasy Plus Micro Kit (Qiagen), according to the manufacturer's instructions, and stored at –80 °C.

**Library preparation and sequencing**

RNA integrity was assessed using a Bioanalyzer (Agilent), and samples with an RNA Integrity Number (RIN) > 7 and RNA input of 60 ng were used for library preparation. Poly(A)-enriched, strand-specific libraries were generated using the NEBNext Ultra II Directional RNA Library Prep Kit (New England Biolabs) by the Molecular Core Facility at CRUK Manchester Institute. Sequencing was performed on an Illumina NovaSeq 6000 platform with an SP flow cell (200-cycle kit), generating 2 × 101 bp paired-end reads at a depth of approximately 20 million reads per sample.

**RNA-seq data processing and analysis**

Raw sequencing reads were assessed using FastQC and FastQ Screen (Babraham Bioinformatics). Adapter sequences were removed, and quality trimming was performed using Trimmomatic v0.39[1]. Reads were aligned to the *Mus musculus* reference genome (GRCm39/mm39) and gene-level counts were obtained using GENCODE M35 annotation and STAR aligner v2.7.7a[2]. Immunoglobulin, mitochondrial, and lncRNA mouse orthologues from https://github.com/BKI-immuno/neoantigen-specific-T-cells-NSCLC were excluded, as were T cell receptor (TCR) genes, ribosomal genes, and haemoglobin genes[3]. Normalisation, principal component analysis, and differential expression analysis were conducted in DESeq2 v1.44.0[4] with default parameters. Differentially expressed genes (DEGs) were defined as those with an adjusted $p$-value *(padj)* $< 0.05$ and a $\log_2$FC $> 1$.

**CIBERSORTx Deconvolution**

Single-cell RNA sequencing (scRNA-seq) data from the ProjecTILs murine reference atlas of tumour-infiltrating T cells[5] was downloaded and imported into R (4.1.3). The Seurat object was split, and 70% of the cells were used to generate a CIBERSORTx reference signature matrix using the cibersortx/fractions Docker image with default settings. This reference matrix was subsequently applied to deconvolute the bulk RNA-seq datasets. To validate the approach, the remaining 30% of the cells were aggregated into pseudo-bulk samples and deconvoluted using the same reference, allowing comparison between predicted and actual cell-type proportions.

**Data visualization**

For heatmap visualisation, normalised gene expression counts were $\log_2$-transformed [$\log_2$(normalised count + 1)] and z-score normalised across each gene (row-wise) to highlight relative expression patterns across conditions. Heatmaps were generated using the ComplexHeatmap R package in R (v4.3.0). Volcano plots were generated by plotting the $\log_2$FC against the $-\log_{10}$ of *padj*. Differential expression data of genes of interest used for heatmap and volcano plots are provided in Supplementary Table S4.

**Gene Set Enrichment Analysis (GSEA)**

The DESeq2 result was filtered to remove genes with p-adjusted NA (genes that did not survive multiple testing correction due to independent filtering). The result was then filtered for duplicate gene symbols (keeping the ones with the greatest basemean) before being ranked by the test statistic (positive to negative). The ranked gene list was analysed using Cluster Profiler GSEA function[6, 7] with settings minGSsize = 15, maxGSSize = 500, and enriched terms were returned with a Benjamini–Hochberg *p*-value cutoff of 1.0 to enable subsequent downstream filtering.

GSEA was performed against the full MSigDB C7: immunological gene signatures, C2: Reactome pathway, and C5: Gene Ontology biological processes collections[8]. A padj value < 0.25 was used as the significance threshold[9].

For visualisation purposes, selected gene sets from the C7 category were annotated as follows: GSE15330 HSC vs Megakaryocyte erythroid progenitor Up: Haematopoietic stem cells (HSC); GSE37301 Multipotent progenitor vs Common lymphoid progenitor Up: Multipotent progenitor cells (MPP); GSE9650 Naive vs Memory CD8 T cell Up: Naïve $CD8^+$ T cells; GSE14699 Naive vs Act CD8 T cell Down: Activated $CD8^+$ T cells; KAECH Day8 Eff vs Memory CD8 T cell Down: Memory $CD8^+$ T cells; GSE15750 Day6 vs Day10 Eff CD8 T cell Up: Proliferative effector $CD8^+$ T cells; GSE9650 Naive vs Exhausted CD8 T cell Down: Exhausted $CD8^+$ T cells.

**Mass cytometry (CyTOF)**

KP475 murine NSCLC tumours were enzymatically dissociated into single-cell suspension as described above. For each condition, a pooled suspension was generated from 4–5 tumours, and $3 \times 10^6$ cells were stained with a panel of metal-conjugated antibodies (1 µl antibody per channel; Supplementary Table S5) in a total of 50 µl Cell staining buffer (Standard Biotools). Viability was assessed by cisplatin (198Pt; Standard Biotools) exclusion. Surface staining was performed after Heparin (Sigma) treatment and Fc blocking (Biolegend). Cells were then fixed and permeabilised using the FOXP3/Transcription Factor Staining Buffer Set (eBioscience), barcoded using palladium 20-plex Barcodes (Standard Biotools), pooled and stained for intracellular antibodies. Samples were fixed overnight in 4% paraformaldehyde (Pierce) /Maxpar PBS (Standard Biotools) at 4 °C and subsequently labelled with Iridium intercalator (125 nM; Standard Biotools) to enable singlet gating based on DNA content.

Barcoded samples were filtered twice through a 70 µm Filcon filter and acquired on a CyTOF XT instrument (Fluidigm) at a rate of ~400 events per second. Data were normalised using EQ Six Element Calibration Beads and debarcoded with CyTOF XT software. Live singlet $CD3^+$ cells were gated using a standard quality control pipeline based on DNA content, EQ bead exclusion, event length, and $cisplatin^-$ signal, performed in FlowJo software. Gated populations were exported as FCS files and analysed in Cytobank using tSNE for dimensionality reduction and FlowSOM for clustering.

**Analysis of clinical data**

To evaluate the clinical relevance of BO-112-induced Tscm enrichment, we generated a $CD8^+$ Tscm gene signature based on differentially expressed genes (DEGs) from RNA sequencing of murine intratumoural $CD8^+$ T cells. Genes enriched in the BO-112 or combination-treated groups were cross-referenced with those previously associated with Tscm $CD8^+$ T cell phenotypes in the literature. The resulting murine gene list was converted to human orthologues using g:Profiler, retaining only 1:1 orthologues for downstream analysis.

Transcriptomic and clinical data from three independent patient cohorts were analysed: TCGA-LUAD (lung adenocarcinoma; n = 522), GSE72094 (lung adenocarcinoma; n = 398), and IMVigor210 (urothelial carcinoma; n = 348). Details of transcriptomic profiling and clinical annotation for these datasets have been previously described[10-12].

Tscm scores were calculated as the median expression of the following signature genes: *SELL*, *LEF1*, *TCF7*, *KLF2*, *CCR7*, *ID3*, *BACH2*, *IL7R*, *FOXO1*, *IKZF1*, *CXCR5*, *SLAMF6*, *BTLA* and *CD27* (Supplementary Table S2). Patients were stratified into "high" and "low" groups based on median dichotomisation (50/50 split). The prognostic value of the Tscm gene signature was assessed using Kaplan–Meier survival analysis and Cox regression (univariable and multivariable). Multivariable analysis was performed only for TCGA-LUAD cohort due to limited clinical metadata in the other datasets. Statistical analyses were conducted in R (v4.3.0) using the packages tibble, survival, and survminer. A $p$-value < 0.05 was considered statistically significant.
